# Shedding light on left hippocampal mGlu5 in Alzheimer’s disease

**DOI:** 10.1101/2025.10.03.680254

**Authors:** Charleine Zussy, Magalie Mathias, Mathieu Vitalis, Xavier Gòmez-Santacana, Cyril Goudet, Catherine Desrumaux, Amadeu Llebaria, Laurent Givalois

## Abstract

Metabotropic glutamate receptor 5 (mGlu5) plays a central role in synaptic plasticity and memory, and has emerged as a potential therapeutic target in Alzheimer’s disease (AD). While asymmetries in hippocampal function have been observed in AD patients, the lateralized contribution of mGlu5 signaling to cognitive decline remains unclear. Here, we show the presence of a physiological left-right asymmetry in mGlu5 expression in the hippocampus of wild-type mice, with higher levels in the left hemisphere, consistent with previous observations. Importantly, we reveal that this asymmetry is lost in J20 AD model mice due to a selective reduction of mGlu5 in the left hippocampus. Then, using the light-controllable negative allosteric modulator Alloswitch-1, we demonstrate that selective inhibition of mGlu5 in the left, but not right, hippocampus is both necessary and sufficient to restore working and short-term memory in J20 mice. This left-specific modulation also reverses downstream pathological signaling, including aberrant Pyk2 and GSK3-β activation and tau hyperphosphorylation, in both hippocampi. Our findings identify a functional lateralization of mGlu5 in hippocampal circuits and highlight the potential of spatially targeted photopharmacology for precise intervention in early AD pathology.

## INTRODUCTION

Alzheimer’s disease (AD) is the most prevalent age-related neurodegenerative disorder, characterized by progressive memory loss, cognitive decline, and behavioral changes. The two hallmark lesions are intracellular neurofibrillary tangles (NFTs), composed of hyperphosphorylated tau, and extracellular β-amyloid (Aβ) plaques (*DeTure and Dickson, 2019; Selkoe, 2001*). However, cognitive impairments in AD correlate more strongly with the levels of soluble oligomeric Aβ (oAβ) than with plaque burden (*Lue et al., 1999; McLean et al., 1999*). Supporting this, transgenic mouse models of AD that overexpress human amyloid precursor protein (hAPP) exhibit deficits in synaptic plasticity and memory prior to the appearance of Aβ plaques (*Holcomb et al., 1998; Oddo et al., 2006; Zussy et al., 2022*). oAβ was shown to impair long-term potentiation (LTP) and enhance long-term depression (LTD) in the hippocampus, two opposing forms of synaptic plasticity that are fundamental to learning and memory, underscoring the synaptotoxic effects of soluble Aβ species and their contribution to cognitive decline in AD (*Shankar et al., 2008*). In this context, the metabotropic glutamate receptor 5 (mGlu5), which is highly expressed in the hippocampus, a region affected early in AD, is critically involved in the regulation of both LTP and LTD, and thus in memory-related processes (*Bikbaev and Manahan-Vaughan, 2017; Hagena and Manahan-Vaughan, 2022; Mukherjee and Manahan-Vaughan, 2013; Ng et al., 2023; Valdivia et al., 2023*). Numerous studies highlight the alteration, contribution and potential therapeutic targeting of mGlu5 in AD (*Abd-Elrahman and Ferguson, 2022; Budgett et al., 2022; Bukke et al., 2020; Kumar et al., 2015*). A recent study has shown that oAβ facilitates mGlu5-dependent LTD through a mechanism involving NMDARs and complement C5aR1 signaling (*Ng et al., 2023*), pointing to aberrant mGlu5 activation as a potential early trigger of synaptic loss. oAβ also hijacks muscarinic receptor-dependent LTD to favor mGlu5-driven depression while inhibiting LTP, a dual effect blocked by preventing Aβ binding to cellular prion protein (PrPᶜ), highlighting a functional interplay between Aβ, PrPᶜ, and mGlu5 in the regulation of synaptic plasticity (*Hu et al., 2014*). Consistently, Aβ from AD brains was found to specifically bind to the N-terminus of PrPᶜ (*Dohler et al., 2014*). This oAβ-PrPᶜ binding was shown to disrupt the PrPᶜ regulatory control over excessive NMDA receptor activity *(You et al., 2012*). Strittmatter’s group also described the existence of an oAβ-PrPᶜ-mGlu5 complex at the synapse, proposing that this interaction leads to aberrant mGlu5 signaling, increased intracellular Ca²⁺ release, and tau hyperphosphorylation underlying neurofibrillary tangle formation (*Abd-Elrahman and Ferguson, 2022; Brody and Strittmatter, 2018; Haas et al., 2016; Laurén et al., 2009; Ng et al., 2023; Salardini et al., 2024; Spurrier et al., 2022; Um et al., 2013*). In addition, oAβ binding to PrPᶜ at the neuronal membrane has been shown to activate the non-receptor tyrosine kinase Fyn, which phosphorylates tau and enhances NMDA receptor signaling, contributing to synaptic dysfunction (*Chin et al., 2005; Crestini et al., 2022; Rajani et al., 2021; Roberson et al., 2011; Um et al., 2012*). Proline-rich tyrosine kinase 2 (Pyk2), a tau kinase encoded by the AD risk gene *PTK2B*, has also been implicated in linking the Aβ-PrPᶜ-mGlu5 complex to tau pathology and neurodegeneration. Pyk2 has been reported to phosphorylate tau directly and to enhance tau hyperphosphorylation indirectly *via* activation of GSK-3β. (*Dourlen et al., 2017; Hartigan et al., 2001; Lambert et al., 2013; Li and Götz, 2018*). While further research is needed to fully characterize the Aβ-PrPᶜ-mGlu5 interaction and signaling, mGlu5 emerges as a critical modulator within this pathological cascade.

A recent report based on electrophysiology experiments showed that oAβ-induced LTD is preferentially enhanced in the left hippocampus through a mGlu5-dependent mechanism (*O’Riordan et al., 2018*). While hippocampal asymmetry, as left-sided atrophy, has been correlated with memory loss in AD (*Fox et al., 1996; Miller et al., 2007; Scahill et al., 2002; Yu et al., 2024*), the synaptic and molecular mechanisms underlying such lateralized vulnerability remain elusive. Although the impact of oAβ on mGlu5 signaling, synaptic plasticity, and memory has been widely investigated, the functional lateralization of hippocampal mGlu5 in AD-related memory deficits remains largely unexplored.

To address this gap, we employed photopharmacology to precisely and region-specifically manipulate mGlu5 activity in awake, behaving mice. This innovative strategy relies on the use of light-regulated compounds (e.g., photoswitchable drugs) to modify the pharmacological properties of target receptors by using specific wavelengths of light to reversibly control drug-receptor interactions. This approach enables the fine spatiotemporal control of drug activity that results in an increased efficacy and selectivity of the drug action (*Gómez-Santacana et al., 2022; Panarello et al., 2022; Velema et al., 2014; Zussy et al., 2018*). Unlike optogenetics, photopharmacology modulates endogenous receptors without the need for genetic modification, offering translational potential for therapeutic development. Alloswitch-1 is a photoswitchable mGlu5 NAM (Negative Allosteric Modulator), which contains an azobenzene-group enabling light-induced configuration changes that result into a reversible loss of affinity for mGlu5. Such photoswitching of mGlu5 activity has been proven both *in vitro* and in different models *in vivo*, making it a very useful tool compound to chart mGlu5 contributions in several neurological diseases (*Gómez-Santacana et al., 2017; Notartomaso et al., 2024; Pittolo et al., 2014; Ricart-Ortega et al., 2020*).

In the present work, we found that wild-type (WT) mice naturally exhibit higher mGlu5 levels in the left hippocampus, a pattern disrupted in the J20 AD model, where mGlu5 expression decreases on the left side, leading to a loss of asymmetry. Then, thanks to Alloswitch-1, we demonstrated that negative modulation of mGlu5 specifically in the left hippocampus was both necessary and sufficient to restore short term memory deficits in AD mice, underscoring the prominent role of this hemisphere in memory recovery. Additionally, we reversed the pathological Aβ-PrPᶜ-mGlu5 signaling cascade, as left-sided modulation of mGlu5 led to reduced Pyk2 and GSK-3β activation, ultimately attenuating tau hyperphosphorylation in the hippocampus of both hemispheres. Our findings reveal a critical role of left hippocampal mGlu5 in AD-related memory dysfunction and highlight photopharmacology as a promising approach for precise therapeutic modulation of pathological signaling pathways.

## MATERIALS AND METHODS

### Animals

Mice were housed in a standard animal facility under a 12 h light/dark cycle at 21 ± 2 °C with *ad libitum* access to food (SAFE Diets, France) and water. All experimental procedures complied with the European Directive 2010/63/EU and national French guidelines for animal welfare. Protocols were approved by the French Ministry of Research and the University of Montpellier Ethics Committee (authorization CEEA-LR-APAFIS#32577). Efforts were made to minimize animal numbers and suffering, and follow the 3Rs rule. The J20 mouse model of AD (*Mucke et al., 2000*) overexpresses human APP carrying the Swedish (K670N/M671L) and Indiana (V717F) mutations under the PDGF-β promoter. Hemizygous J20 mice were obtained from The Jackson Laboratory and bred on a C57BL/6J background (>10 backcrosses) under a material transfer agreement with the Gladstone Institute (#UM140255-01). Genotyping was performed as described previously (*Mansuy et al., 2018*). Both sexes were used and pooled, as no sex differences were observed. Naive mice received no pharmacological treatment but were handled in the same manner as treated animals. Vehicle-treated mice received the appropriate vehicle (intraperitoneally (i.p.) or intrahippocampal) and served as negative controls.

### Drugs

The photoswitchable negative allosteric modulator of mGlu5, Alloswitch-1, was synthesized in the MCS laboratory (Barcelona, Spain) following previously published protocols (*Gómez-Santacana et al., 2017; Pittolo et al., 2014*).

### Intraperitoneal Drug Administration

For dose-response experiments, Alloswitch-1 was injected i.p. at 10 or 20 mg/kg (5% DMSO, 5% Tween 20 in saline) 40 minutes before behavioral testing (Suppl. Fig.2A-B).

### Stereotaxic Surgery and Optical Fiber/Cannula Implantation

Mice were anesthetized with ketamine/xylazine and placed in a stereotaxic frame. Guide cannulas or optrodes were implanted targeting the CA1 region of the dorsal hippocampus (A/P: −2.0 mm, M/L: ±1.8 mm, D/V: −1.25 mm) using flat skull coordinates (*De Bundel et al., 2016; Iaccarino et al., 2016*). Cannula (26 gauge, 2.50 mm, Plastics One) or optrodes (Doric Lenses) were secured with anchor screws and dental acrylic (AgnTho’s), and protected postoperatively with dummy cannula. A recovery period of at least 7 days was allowed before testing, as previously reported (*Zussy et al., 2018*).

### In Vivo Photopharmacological Experiments

For experiments involving the inactivation of Alloswitch-1 in the left or right CA1, mice were implanted with an optical fiber targeting the corresponding hemisphere and allowed to recover for one week. Alloswitch-1 was administered i.p. at an effective dose of 20 mg/kg (as previously determined; Suppl. Fig.2B), 40 minutes before the behavioral task. Optical stimulation was delivered using a compact LED connected to a 200 µm-diameter rotating optical fiber (NA = 0.48), controlled via LED driver software (Doric Lenses). Violet light (λ = 385 nm; 50 ms pulses at 10 Hz, 1-2 mW) was applied starting 20 minutes before the task and maintained throughout the 8-minute Y-maze session (Fig.2A).

**FIGURE 1.**
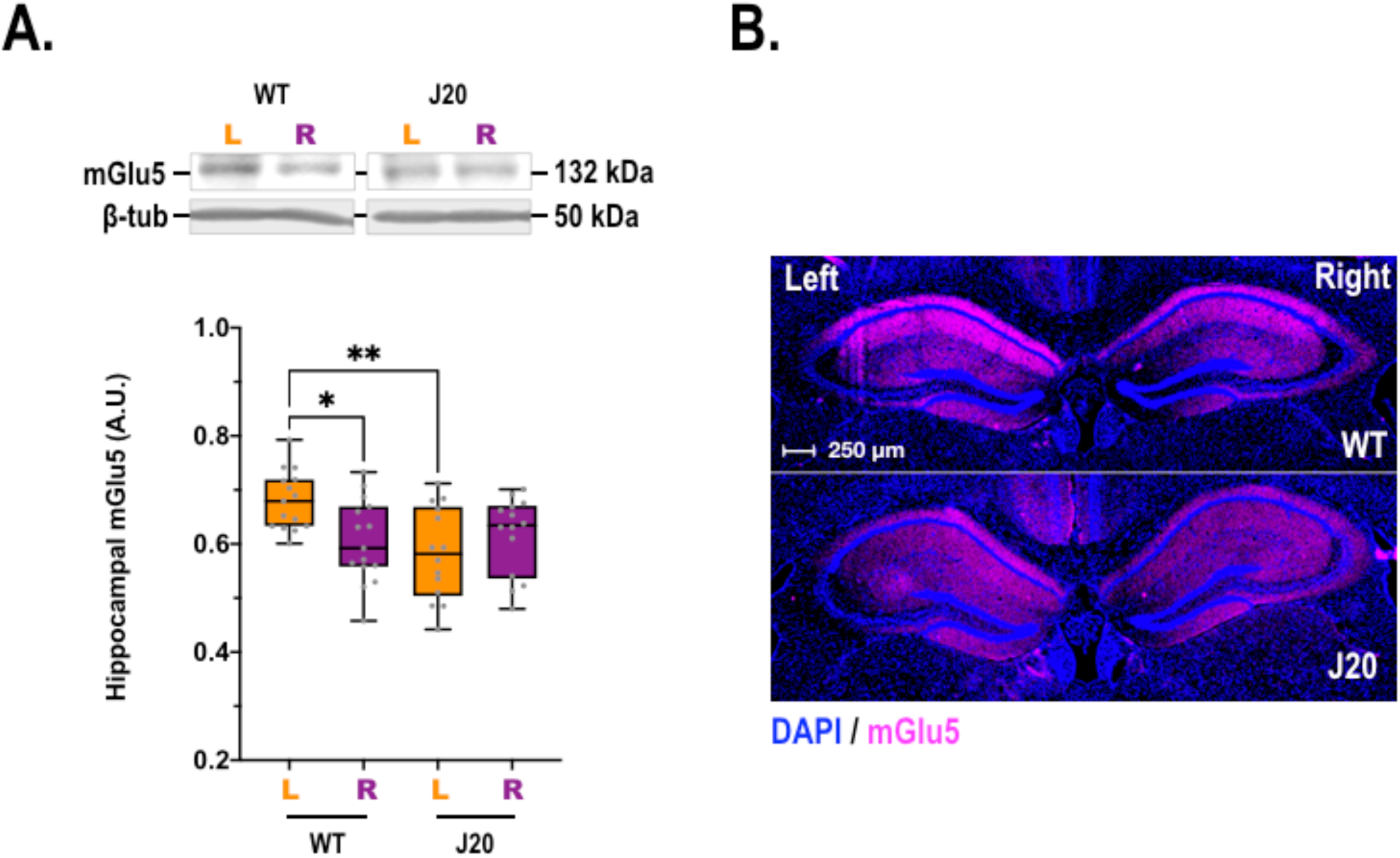
**A.** mGlu5 protein levels (132 kDa) were measured in the **left** (L) and **right** (R) hippocampus of WT and J20 mice by Western blot and normalized to β-tubulin (β-tub, 50 kDa). Data are expressed as the mGlu5/β-tub ratio. Two-way ANOVA followed by Tukey’s post hoc test: genotype, F_1,46_= 4.50, p = 0.0392; lateralization, F_1,46_= 1.26, (ns); interaction, F_1,46_= 9.30, p = 0.0038. * p< 0.05 and ** p< 0.01 vs. WT-L. Data are presented as box-and-whisker plots with Min to Max and Median, gray dots represent individual animals. **B.** Representative immunohistochemical images of hippocampal sections (12 µm) from WT and J20 mice stained for mGlu5 (magenta) and counterstained with DAPI (blue). Scale bars: 250 µm.

**FIGURE 2.**
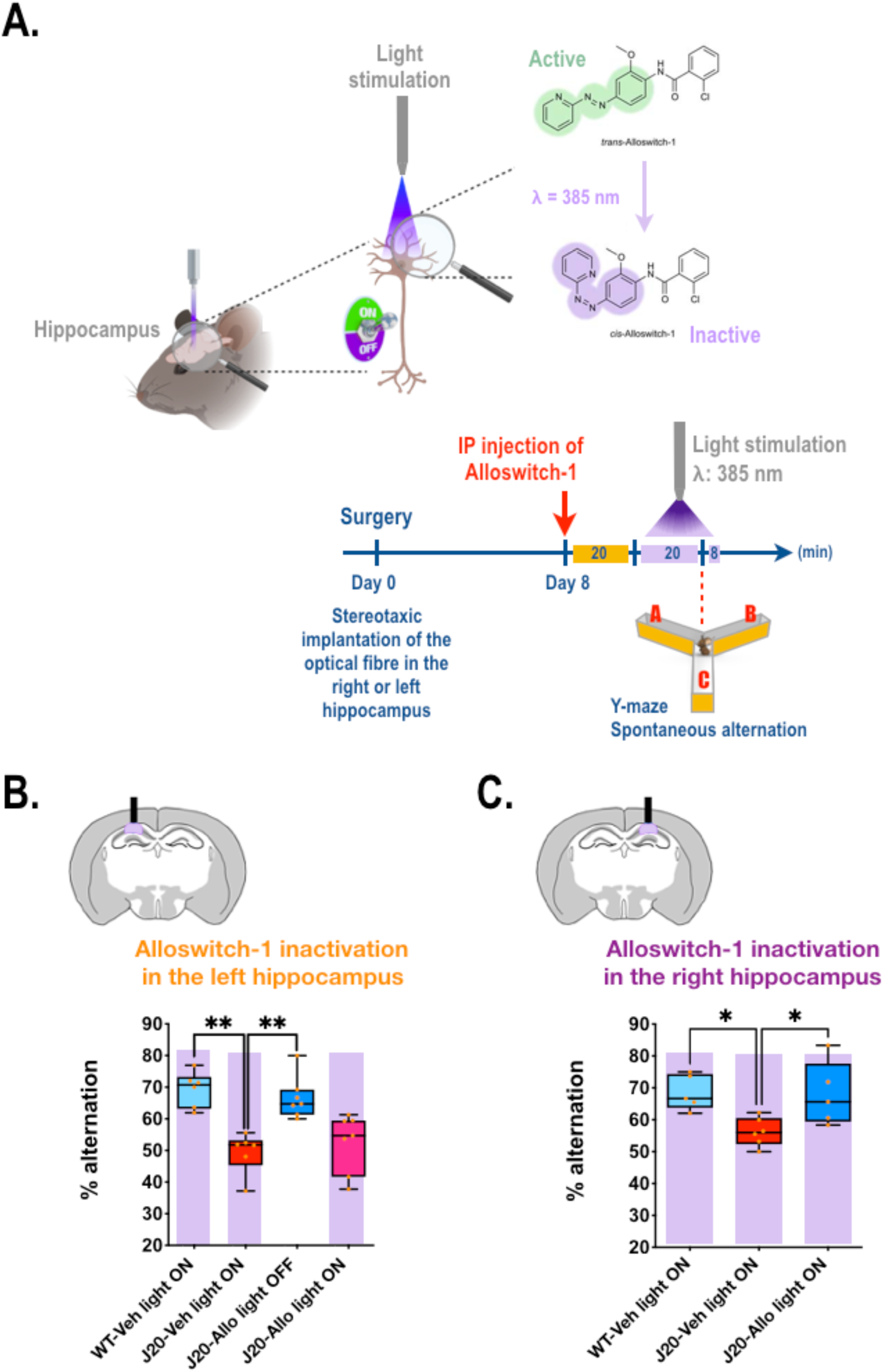
**A.** *Inactivation protocol for Alloswitch-1*. One week after stereotaxic implantation of an optical fiber targeting either the left or right hippocampus, mice received an intraperitoneal (i.p.) injection of Alloswitch-1 (Allo, 20 mg/kg) or vehicle (Veh). Violet light stimulation (385 nm) was applied for 20 minutes prior to, and during, the 8-minute Y-maze test to locally inactivate Alloswitch-1 in a hemisphere-specific manner. Spontaneous alternation behavior was used as a measure of spatial working memory, calculated as: (number of spontaneous alternations / [total arm entries – 2]) × 100. **B-C.** Working memory performance in the Y-maze following Alloswitch-1 inactivation in the **left** (**B**) or **right** (**C**) hippocampus. One-way ANOVA followed by Dunnett’s multiple comparisons: **left** (**B**): F_3,22_=12.0 (p< 0.0001). ** p< 0.01 *vs.* J20-Veh light ON group; **right** (**C**): F_2,13_=5.65 (p= 0.0171). * p< 0.05 *vs.* J20-Veh light ON group. Data are presented as box-and-whisker plots (min to max, with median), orange dots represent individual animals. Note that a part of this figure was created in BioRender.

### Intrahippocampal Drug Infusion

For experiments involving direct hippocampal administration, mice were implanted with a guide cannula targeting the left or right CA1 and allowed to recover for one week. Consistent with previous protocols, Alloswitch-1 (300 nM in 0.003% DMSO in PBS) or vehicle was infused (2 µl at 0.5 µl/min) into the corresponding hemisphere (*Gómez-Santacana et al., 2017; Ricart-Ortega et al., 2020*). The injector was left in place for 6 minutes post-infusion to ensure proper diffusion and prevent reflux. Working memory and spatial novelty preference were assessed in the Y-maze 20 and 15 minutes, respectively, after the start of the infusion (Fig.3A,C).

**FIGURE 3.**
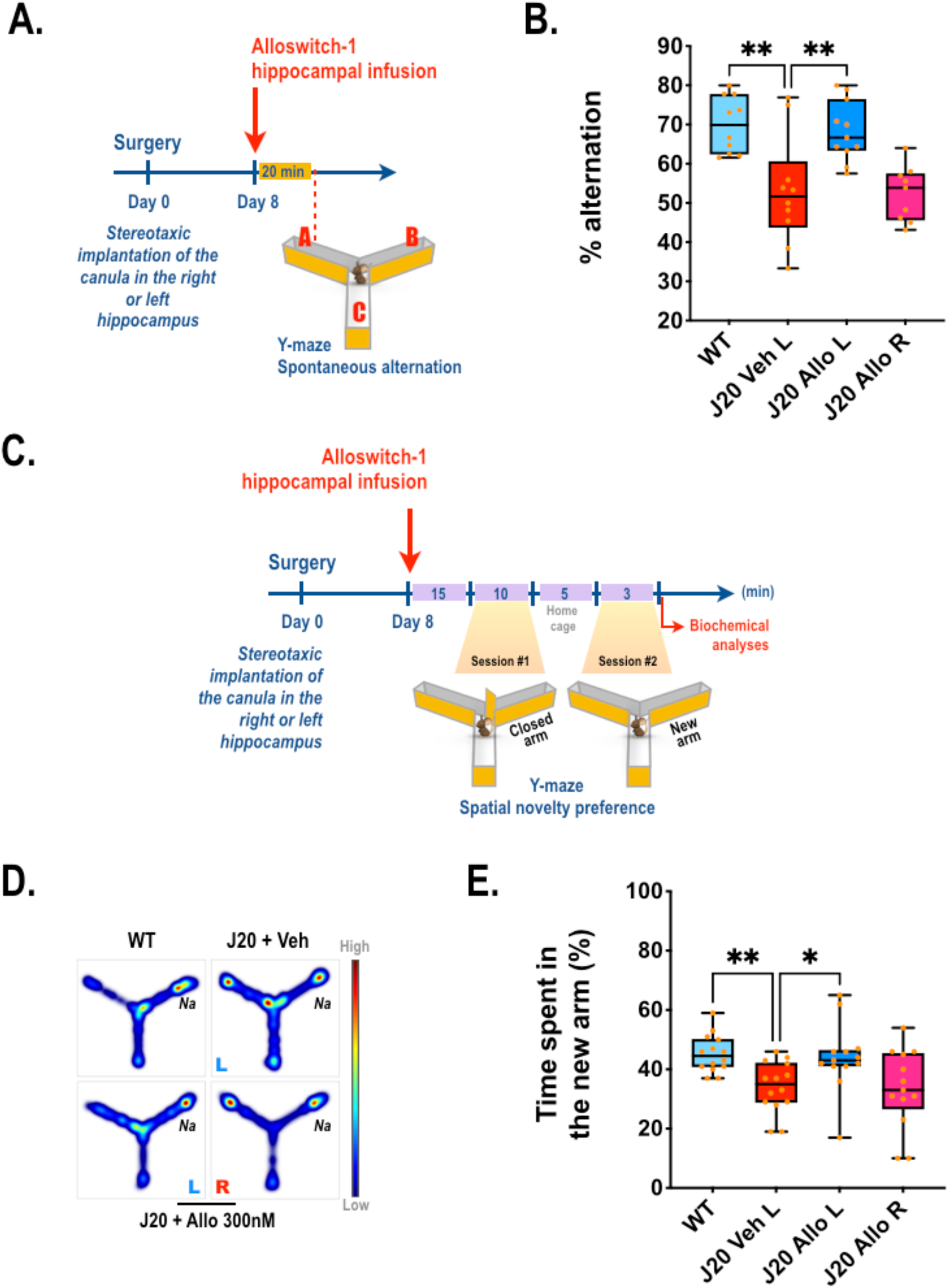
**A,C.** *Intrahippocampal infusion protocol.* One week after stereotaxic implantation of a guide cannula, Alloswitch-1 (Allo, 300 nM) or vehicle (Veh) was infused into the left or right hippocampus 20 min before the Y-maze working memory test (**A**) or 15 min before the spatial novelty preference test (**C**). **B.** Effect of Alloswitch-1 infusion (300 nM) into the **left** (L) or **right** (R) hippocampus on working memory performance. Data are expressed as percentage of spontaneous alternations calculated as (spontaneous alternations / [number of arm entries − 2]) × 100. One-way ANOVA followed by Dunnett’s multiple comparisons: F_3,36_= 10.1 (*p* < 0.0001). ** p< 0.01 *vs*. J20-Veh L group. **D-E.** Effect of Alloswitch-1 infusion (300 nM) into the **left** (L) or **right** (R) hippocampus on short-term spatial memory. **D.** Representative heatmaps for each group. **E.** Percentage of time spent in the novel arm. One-way ANOVA followed by Dunnett’s multiple comparisons: F_3,36_ = 10.09 (p < 0.001). * p< 0.05 and ** p< 0.01 *vs*. J20-Veh L group. All data are presented as box-and-whisker plots (Min to Max, with Median), orange dots represent individual animals. Note that a part of this figure was created in BioRender.

### Y-Maze Behavioral Testing

#### Spontaneous Alternation Task (Working Memory)

This task assesses spatial working memory based on rodents’ innate preference for exploring novel environments. The Y-maze consisted of three identical arms (35 cm long × 6 cm wide, with 15 cm high walls) arranged at 120° angles. As previously described (*Canet et al., 2025*), mice were placed at the end of one arm and allowed to freely explore all three arms for 8 minutes. Spontaneous alternation behavior was calculated as: (Number of alternations / [number of arm entries − 2]) × 100. Maze cleaning was performed between animals using 50% ethanol. Tracking was conducted using EthoVisionXT (Noldus).

### Spatial Novelty Preference Task (Short-Term Memory)

The test was adapted from Shipton et al., 2014 (*Shipton et al., 2014*). During the exploration phase (10 min), access to the novel arm was blocked. After a 5 min delay, the block was removed and mice were allowed 3 min to explore all arms. Time spent in each arm was recorded from the moment the mouse exited the start arm. To minimize olfactory bias, the novel arm was wiped with a tissue previously rubbed along the explored arms. The percentage of time spent in the novel arm was used as an index of spatial short-term memory. Maze cleaning and tracking followed the same procedure as above.

### Soluble Aβ_1-42_ assay

Mice were euthanized by decapitation, hippocampi were dissected, frozen in liquid nitrogen, and stored at −20 °C. As previously reported, frozen hippocampi were sonicated in a lysis buffer (140 mM KCl, 10 mM Na_2_HPO_4_, 1.7 mM KH_2_PO_4_, 1 mM EDTA) containing a cocktail of protease and phosphatase inhibitors (Roche Diagnostics GmbH, Germany) (*Mansuy et al., 2018*). The homogenate was then centrifuged at 4°C (16000 g, 25 min) and the supernatant containing soluble fraction was collected. Soluble human amyloid-β(1-42) (Aβ_1-42_) peptide was quantified using an ELISA kit according to the manufacturer’s protocol (Human Aβ42 Ultrasensitive ELISA Kit, Invitrogen). The results were normalized to protein concentrations, determined using the bicinchoninic acid method (BCA kit, ThermoFisher Scientific).

### Western blot analyses

Immediately after the spatial novelty preference task, mice were euthanized by decapitation. Hippocampi were dissected, frozen in liquid nitrogen, and stored at −20 °C. As previously reported, tissues were homogenized by sonication in 3% SDS containing protease and phosphatase inhibitors (Roche Diagnostics GmbH) (*Zussy et al., 2022*). Protein concentration was determined using a BCA assay (ThermoFisher Scientific). Thirty micrograms of protein were loaded per lane, separated by SDS-PAGE (12%), and transferred onto nitrocellulose membranes. After overnight incubation with primary antibodies at 4 °C, membranes were washed and incubated with HRP-conjugated secondary antibodies (Sigma-Aldrich) for 2 hours. Signal was revealed using ECL (Luminata Crescendo, Millipore) and quantified with ImageJ software (NIH). β-Tubulin was used as a loading control for mGlu5, while total Pyk2, Fyn, and tau served for their respective phosphorylated forms.

The following antibodies were used: rabbit anti-mGlu5 (1/500, Millipore), rabbit anti-phospho-Pyk2 (Tyr402, 1 /500 Cell Signaling), mouse anti-Pyk2 (clone 5E2, 1/1000, Cell Signaling), mouse anti-phospho-GSK-3β (Tyr216, 1/2000, BD biosciences), mouse anti-GSK-3β (1/2000, BD biosciences), rabbit anti-phospho-Fyn (Tyr416, 1/500, Cell Signaling), rabbit anti-Fyn (1/500, Cell Signaling), mouse anti-phospho-tau (Thr231, clone AT180, 1/1000, Thermofisher), rabbit anti-tau total (1/10000, Dako Cytomation), mouse anti-β-tubulin (1/7500, Sigma Aldrich).

### Fluorescent immunohistochemistry

Animals were anesthetized (Exagon, 40 mg/kg, i.p.) and perfused with PBS followed by 4% PFA. Brains were post-fixed in 4% PFA for 48 h at 4 °C, then cryoprotected in 30% sucrose. As previously reported, brains were embedded in OCT (Tissue-Tek), frozen in dry ice–chilled acetone, and sectioned (*Canet et al., 2025; Zussy et al., 2018*). Twelve-micrometer coronal brain sections were obtained using a cryostat (Leica CM3050-S). Slides were incubated overnight at room temperature with rabbit anti-mGlu5 (1:500, Millipore), followed by Cy5-conjugated anti-rabbit secondary antibody (1:2000, Invitrogen). Nuclei were counterstained with DAPI. Images were acquired using a Leica DM4B microscope with Thunder Imager (Leica microsystem).

### Exclusion criteria and group analysis

Animals were excluded based on predefined criteria: (1) weight loss or illness post-surgery; (2) blocked or dislodged cannula or ineffective light delivery; (3) misplacement of implants; (4) abnormal behavior (e.g., excessive aggression). Experiments could not be fully blinded due to factors such as compound color, transgenic phenotype, and visible light delivery; however, behavioral scoring was automated using the EthoVision XT system to minimize bias.

### Statistical analysis

Data are presented as box-and-whisker plots (min to max, with median) and each data point on the graphs represents an individual animal. Normality was assessed using the Kolmogorov–Smirnov test. One- or two-way ANOVA with Dunnett’s or Tukey’s post hoc tests, respectively, was applied to normally distributed data. A one-sample t-test was used to compare the sample mean to the theoretical value of 1.0. Statistical analyses were conducted using GraphPad Prism 9.0. A p-value < 0.05 was considered statistically significant. Detailed statistical information is provided in the figure legends. The number of animals and statistical details are indicated in the figure legends. Statistical power analysis was calculated using G*Power.

## RESULTS

### Asymmetric hippocampal mGlu5 expression is lost in a mouse model of AD

To investigate the contribution of hippocampal mGlu5 to memory deficits in J20 mice, we first assessed mGlu5 expression using western blotting and immunofluorescence. Quantitative western blot analysis revealed three key findings: (1) in WT mice, mGlu5 expression is higher in the left than in the right hippocampus; (2) in J20 mice, mGlu5 expression is selectively reduced in the left hippocampus compared to WT controls; and (3) no interhemispheric difference in mGlu5 levels is observed in J20 mice (Fig.1A). These results were corroborated by immunofluorescent staining, which clearly illustrated the loss of left-dominant mGlu5 expression in J20 mice. Notably, changes in mGlu5 expression were most prominent in the CA1 region of the hippocampus (Fig.1B). Additionally, we observed a trend toward increased levels of soluble Aβ_1–42_ in the left hippocampus (Supplementary Fig.1A), suggesting a possible link between reduced mGlu5 expression and local Aβ accumulation.

### Alloswitch-1 restores memory in J20 mice through selective inhibition of left hippocampal mGlu5

Alloswitch-1 is a photoswitchable NAM of mGlu5 that is active in its thermodynamically stable trans-isomer, which is produced either upon green light illumination or by thermal relaxation in the dark (*Pittolo et al., 2014*) (Fig.2A). Given that mGlu5 NAMs have shown cognitive benefits in multiple AD models (*Abd-Elrahman and Ferguson, 2022*), we first evaluated Alloswitch-1 in the dark as a conventional mGlu5 NAM by i.p. injection in J20 mice and assessed working memory using the Y-maze spontaneous alternation test. In this task, working memory performance is indicated by the percentage of spontaneous alternations, sequential entries into the three arms of the maze, during an 8-minute free exploration period. J20 mice treated with vehicle exhibited reduced alternation compared to WT controls, consistent with impaired working memory, as previously reported (*Canet et al., 2025*). As expected, administration of Alloswitch-1 (20 mg/kg, i.p.) restored performance to WT levels, confirming the therapeutic potential of mGlu5 inhibition in this model (Supplementary Fig.2A-B). To dissect the hemispheric contribution of mGlu5 to memory impairment, we next exploited the light-sensitive properties of Alloswitch-1. Upon illumination with violet light (λ = 385 nm), Alloswitch-1 undergoes isomerization to its cis-form, rendering it inactive (Fig.2A). We implanted an optrode above the left or right CA1 region of the hippocampus to allow localized photo-inactivation of the drug. Following recovery, J20 mice received systemic Alloswitch-1 (20 mg/kg, i.p.) or vehicle, and violet light was applied unilaterally to selectively inactivate the drug in either hemisphere. Behavioral testing in the Y-maze revealed that memory rescue by Alloswitch-1 was completely abolished when the drug was inactivated in the left hippocampus (Fig.2B), whereas inactivation in the right hippocampus had no impact on the drug’s efficacy (Fig.2C). Neither violet light stimulation alone nor vehicle injection affected performance. Importantly, since memory performance in J20 mice following Alloswitch-1 injection did not differ with or without the implanted optrode (Supp Fig.2C), we omitted the right hippocampus "Allo light OFF" control group to reduce animal use in accordance with ethical guidelines.

As left hippocampal mGlu5 appeared to play a dominant role in memory processing, we next examined whether local administration of Alloswitch-1 into this region could rescue cognitive deficits in J20 mice (Fig.3A). To reduce the number of animals used, we selected a concentration of 300 nM based on prior *in vivo* studies reporting behavioral efficacy at this dose (*Gómez-Santacana et al., 2017; Ricart-Ortega et al., 2020*). Local infusion of Alloswitch-1 into the left hippocampus was sufficient to restore working memory in J20 mice, whereas infusion into the right hippocampus had no effect. As expected, vehicle infusion into the left hippocampus failed to improve performance (Fig.3B). To complement the spontaneous alternation Y-maze test, which primarily assesses working memory by relying on the mouse’s tendency to alternate arm choices based on recent entries (*Hughes, 2004*), we employed the spatial novelty preference Y-maze task. This task evaluates short-term memory by measuring the animal’s ability to recognize recently explored spatial contexts and its preference for novelty (*Hughes, 2004; Shipton et al., 2014*). The test was performed following unilateral hippocampal infusion of Alloswitch-1 (Fig.3C). Consistent with results from the spontaneous alternation task, only mice receiving Alloswitch-1 infusion into the left hippocampus exhibited restored short-term memory in the spatial novelty preference task (Fig.3D-E). Importantly, despite the reduced mGlu5 expression observed in the left hippocampus of J20 mice (Fig.1), the receptor remained responsive to Alloswitch-1 treatment (Figs 2 & 3). Together, these results highlight the preferential involvement of left hippocampal mGlu5 in AD-related spatial memory impairment.

### Left hippocampal mGlu5 inhibition reverses bilateral signaling dysfunction in AD

As the Aβ-PrPᶜ-mGlu5 signaling complex has been reported to promote downstream neurotoxic pathways, including activation of Fyn, Pyk2 and GSK-3β kinases that leads to tau hyperphosphorylation, we next measured the levels of these key molecular markers in the hippocampus of J20 mice following Alloswitch-1 infusion, to determine whether mGlu5 modulation could mitigate these pathological effects. We previously investigated tau phosphorylation in J20 mice. Although this model is primarily used to study amyloid pathology, we observed in the hippocampus of 9-month-old mice, a strong hyperphosphorylation of tau at several AD-relevant epitopes (i.e. AT180, AT8, AT270, and CP13), increased activation of related kinases (i.e. GSK-3β, Cdk5, p25/35) and elevated Fyn levels (*Canet et al., 2025*).

We first measured mGlu5 expression levels in the left and right hippocampus to determine whether the treatment could restore the asymmetry observed in WT mice. Alloswitch-1 treatment had no detectable effect on mGlu5 expression levels, regardless of whether it was infused into the left or right hippocampus (Supp Fig.3A-D). It is possible that a single administration of Alloswitch-1 is insufficient to induce detectable changes in mGlu5 expression levels. However, the acute administration of Alloswitch-1 was sufficient to normalize pathological modifications of kinases as Fyn, Pyk2 and GSK-3β (Supp Fig.3 & Fig.4). Phosphorylated Pyk2 levels were elevated in J20 mice that received vehicle infusion, whereas Alloswitch-1 infusion into the left hippocampus restored their phosphorylation to levels comparable to those observed in WT mice. Remarkably, left hippocampal infusion of Alloswitch-1 also restored phosphorylated Pyk2 levels in the right hippocampus. In line with the lateralized effect of Alloswitch-1 on memory rescue, infusion into the right hippocampus failed to reduce Pyk2 phosphorylation levels in J20 in either hemisphere (Fig.4A-D). Phosphorylated (Tyr216) GSK-3β levels were selectively increased in the left hippocampus of J20 mice infused with vehicle (Fig.4E-F, Supp Fig.1B). Infusion of Alloswitch-1 into the left hippocampus normalized GSK-3β phosphorylation to levels observed in WT mice. By contrast, its phosphorylation levels in the right hippocampus remained unchanged across all groups. Additionally, infusion of Alloswitch-1 into the right hippocampus had no detectable effect. (Fig.4E-F). Fyn activation, which was reduced in J20 mice receiving vehicle infusion, was similarly bilaterally restored to WT levels following left hippocampal administration of Alloswitch-1 (Supp Fig.3E-F). Fyn, Pyk2, and GSK-3β are known to act upstream of tau pathology. As recent studies implicate Pyk2 activation in promoting tau phosphorylation, either directly *(Li and Götz, 2018*) or *via* activation of GSK-3β (*Hartigan et al., 2001*) downstream of Aβ-PrPᶜ-mGlu5 signaling, we next examined whether Alloswitch-1 treatment could modulate tau hyperphosphorylation in J20 mice. Among the epitopes previously found to be strongly hyperphosphorylated in this model (*Canet et al., 2025*), we focused on AT180 (T231/S235), a clinically relevant site whose phosphorylation disrupts tau binding to microtubules and promotes early aggregation events in human AD pathology (*Sengupta et al., 1998*). Consistent with its bilateral effects on kinases activation, infusion of Alloswitch-1 into the left hippocampus of J20 mice also resulted in a marked reduction of tau hyperphosphorylation in both the left and right hippocampi, whereas infusion into the right hippocampus had no detectable effect (Fig.4G-H). Strikingly, this reveals that a highly localized intervention in the left hippocampus may engage interhemispheric mechanisms sufficient to elicit broader molecular rescue. Such spatially restricted yet widespread effects highlight a potentially powerful therapeutic entry point in early AD.

**FIGURE 4.**
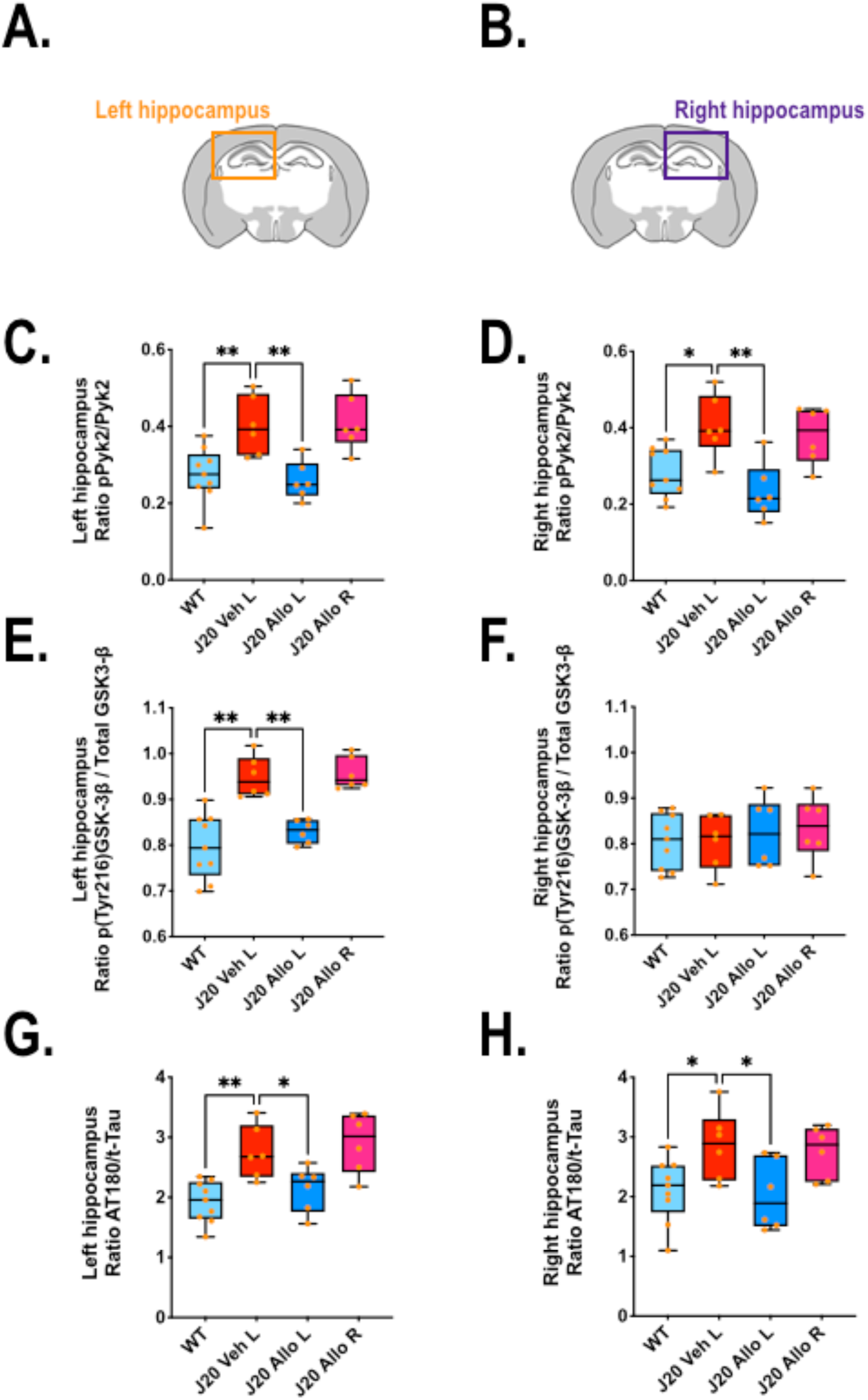
Effect of Alloswitch-1 infusion (300 nM) into the **left** (L) or **right** (R) hippocampus on activation of Pyk2 and GSK-3β kinases and tau hyperphosphorylation. For experimental protocol see Fig.3C. Hippocampal variations of Phospho-Pyk2 protein levels (116 kDa) in the **left** (**A,C**) and **right** (**B,D**) hippocampus were measured by Western blot and normalized to total Pyk2 levels (116 kDa). Data are expressed as the Phospho-Pyk2/Pyk2 ratio. One-way ANOVA followed by Dunnett’s multiple comparisons: **C**: F_3,23_ = 9.04 (p < 0.001); **D**: F_3,23_ = 7.68 (p < 0.001). * p< 0.05 and ** p< 0.01 *vs*. J20-Veh L group. Phospho-GSK-3β protein levels (46 kDa) in the **left** (**A**,**E**) and **right** (**B,F**) hippocampus, normalized to total GSK-3β levels (46 kDa). Data are expressed as the Phospho-GSK-3β / total GSK-3β ratio. One-way ANOVA followed by Dunnett’s multiple comparisons: **E**: F_3,23_ = 18.1 (p < 0.0001); **F**: F_3,23_ = 0.27 (ns). * p< 0.05 and ** p< 0.01 *vs*. J20-Veh L group. Phospho-tau protein levels (AT180, 60 kDa) in the **left** (**A,G**) and **right** (**B,H**) hippocampus, normalized to total tau (t-Tau) levels (60 kDa). Data are expressed as the Phospho-tau (AT180)/t-Tau ratio. One-way ANOVA followed by Dunnett’s multiple comparisons: **G**: F_3,23_ = 9.32 (p < 0.001); **H**: F_3,23_ = 4.16 (p < 0.05). * p< 0.05 and ** p< 0.01 *vs*. J20-Veh L group. All data are presented as box-and-whisker plots (Min to Max, with Median), orange dots represent individual animals. Note that a part of this figure was created in BioRender.

**FIGURE 5.**
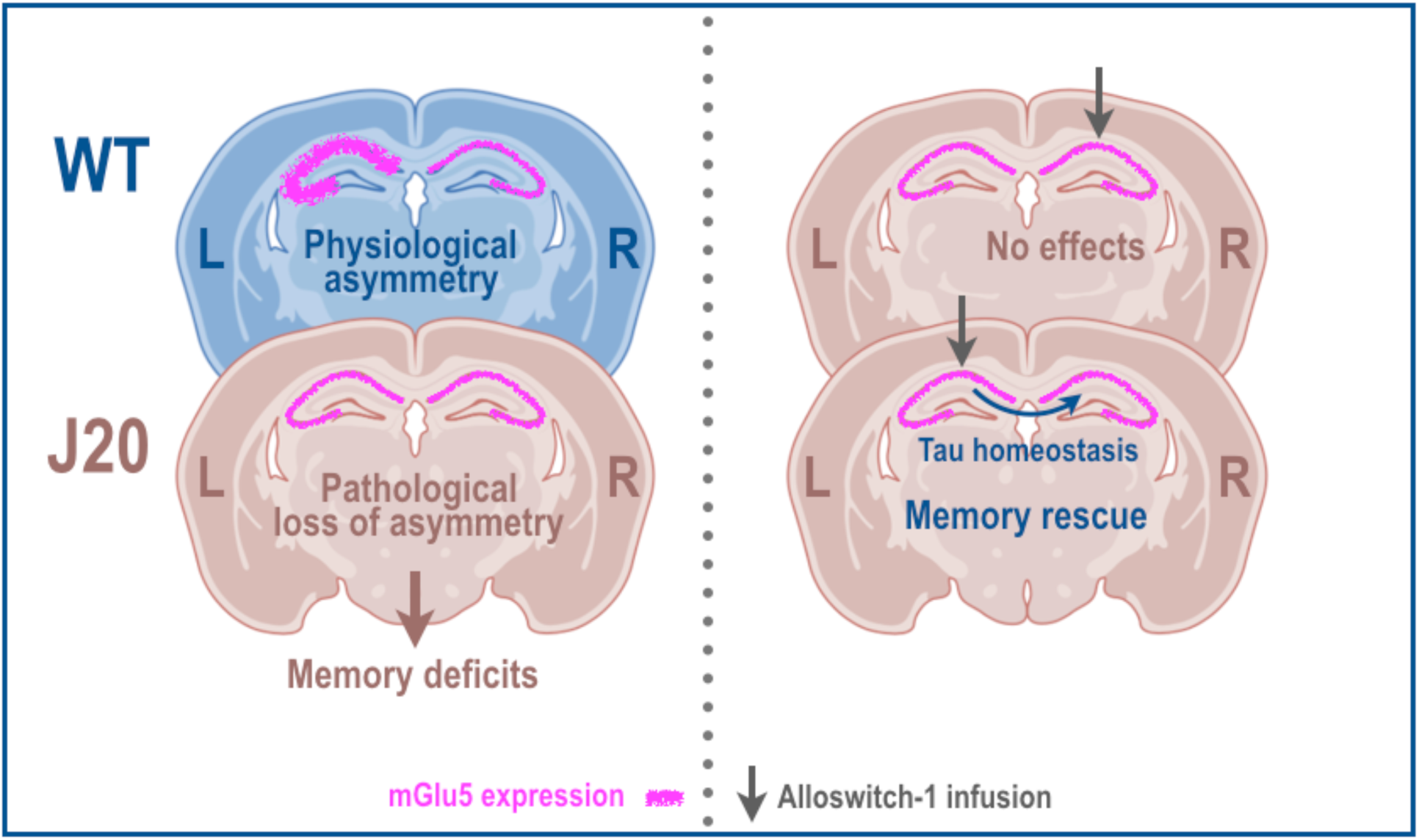
Disrupted left hippocampal mGlu5 asymmetry and functional restoration in J20 mice. Left panel : Schematic coronal sections of the hippocampus in WT (blue) and J20 mice (brown). WT mice show higher mGlu5 expression (pink line) in the left hippocampus, which is lost in J20 mice and associated with memory deficits. Right panel : Infusion of Alloswitch-1 in J20 mice (gray arrow). Infusion into the right hippocampus has no effect on memory, whereas left hippocampal infusion rescues memory performance. Notably, left-sided treatment also normalizes tau pathology in both hemispheres (blue arrow), indicating interhemispheric propagation of therapeutic effects. Note that a part of this figure was created in BioRender.

## DISCUSSION

mGlu5 has emerged as a key molecular player in AD, with substantial evidence supporting its pathological involvement and therapeutic relevance (*Abd-Elrahman and Ferguson, 2022; Budgett et al., 2022; Bukke et al., 2020; Kumar et al., 2015*). However, the potential lateralization of mGlu5 signaling in AD-related memory deficits had not been directly addressed. Here, we confirm that WT mice naturally display asymmetric mGlu5 expression, with higher levels in the left hippocampus. This physiological asymmetry is lost in J20 mice, due to a selective downregulation of mGlu5 in the left hemisphere.

The preferential decrease of mGlu5 in the left hippocampus during disease progression raises important questions about the functional state of the remaining receptors in this region. Notably, despite this left-sided reduction in mGlu5 expression, further pharmacological inhibition of mGlu5 activity in the left hippocampus effectively rescues short-term memory deficits in J20 mice. Although this combined downregulation and inhibition may appear paradoxical, it likely reflects the complex functional dysregulation of mGlu5 signaling in the AD brain. The mGlu5 downregulation may initially represent a compensatory mechanism to counteract glutamate excitotoxicity, a well-established feature of AD. However, as the disease progresses, pathological processes, including altered receptor coupling, aberrant trafficking, or pathological interactions with Aβ oligomers, can convert this adaptive adjustment into maladaptive signaling, rendering the remaining receptors hyperactive and contributing to synaptic dysfunction (*Abd-Elrahman and Ferguson, 2022; Budgett et al., 2022; Bukke et al., 2020; Renner et al., 2010; Um et al., 2013*). Pharmacological blockade may thus restore balanced receptor signaling and synaptic function, improving memory performance despite the lower receptor levels.

Alterations in mGlu5 expression or availability have been documented in both patients and animal models, though findings remain heterogeneous. Recent PET imaging studies in patients have reported altered mGlu5 availability and binding in the AD brain. However, the findings are heterogeneous: one study reported increased mGlu5 binding in the cortex and hippocampus in severe AD cases (*Müller Herde et al., 2019*), while another showed decreased binding specifically in the hippocampus during early stages of the disease (*Mecca et al., 2020*). Additionally, a study in stage-matched AD patients revealed decreased binding of an mGlu5-selective PET tracer in both the hippocampus and the amygdala (*Treyer et al., 2020*). These discrepancies likely reflect region- and stage-specific dynamics, which are mirrored in transgenic models. For example, 5xFAD mice exhibit reduced hippocampal mGlu5 binding at 10 months (*Lee et al., 2018*), whereas Tg-ArcSwe mice show stable or slightly elevated expression across ages (*Fang et al., 2017*). In APP/PS1 mice, findings are also variable: Valdivia et al. (2023) reported no significant changes in hippocampal mGlu5 levels at 8 months, while Privitera et al. (2022) observed a biphasic pattern, with increased expression at 7 months followed by marked downregulation by 13 months (*Privitera et al., 2022; Valdivia et al., 2023*). Collectively, these data support a temporally dynamic regulation of mGlu5 in AD, shaped by genetic background, disease stage, and brain region.

In our study, the reduction of mGlu5 expression was confined to the left hippocampus of 7-month-old J20 mice, corresponding to a moderate stage of disease progression. This disruption of hemispheric asymmetry may reflect an early breakdown of lateralized memory networks. In patients, early structural and functional alterations, including left-predominant cortical atrophy and Aβ accumulation, have been linked to cognitive decline (*Derflinger et al., 2011; Frings et al., 2015; Miller et al., 2007; Tyrer et al., 2020; Weise et al., 2018; Yu et al., 2024b*), suggesting that left-hemispheric vulnerability may be a generalizable feature of AD. Interestingly, this decline is associated by a compensatory recruitment of the right hemisphere (*Tyrer et al., 2020*), potentially reflecting a broader collapse of hemispheric lateralization as the disease advances.

Such pathological alterations may arise from a disruption of normally lateralized hippocampal circuits. In healthy rodents, the hippocampus exhibits intrinsic hemispheric asymmetries at both molecular and functional levels. Synapses in CA1 receiving input from the left CA3 show enrichment in GluN2B-containing NMDA receptors, whereas those receiving right CA3 input are larger and enriched in GluA1-containing AMPA receptors (*Kawakami et al., 2003; Kitanishi et al., 2017; Shinohara et al., 2008*). mGlu5 expression also follows this asymmetric pattern, with higher levels reported in the left CA1 postsynaptic density *(Shinohara and Hirase, 2009*), consistent with our findings in WT animals.

This molecular asymmetry has functional consequences. Using optogenetic approaches, Kohl et al. (2011) showed that selective stimulation of left CA3 projections leads to stronger LTP in CA1 compared to right-sided input (*Kohl et al., 2011*). In a subsequent study, Shipton et al. (2014) demonstrated that unilateral silencing of either left or right CA3 impairs short-term memory, indicating that both hemispheres contribute to this process. However, only silencing of the left CA3 disrupted long-term spatial memory, highlighting a lateralized contribution to memory consolidation (*Shipton et al., 2014*). Converging evidence from human intracranial recordings similarly points to hemispheric lateralization, with the left hippocampus supporting spatial memory and the right hippocampus navigation (*Miller et al., 2018*).

Importantly, mGlu5 lateralization is not restricted to the hippocampus. Kolber et al. (2010) reported that differential expression and activation of mGlu5 in the left versus right amygdala produced distinct behavioral responses to pain stimuli, pointing to a broader principle of hemisphere-specific mGlu5 function across limbic circuits. These findings reinforce the idea that mGlu5 signaling is subject to functional lateralization in multiple brain regions and is behaviorally relevant (*Kolber et al., 2010*).

Our study expands this framework by demonstrating that the rescue of spatial short-term memory in J20 mice specifically depends on mGlu5 activity in the left CA1. Inactivation of Alloswitch-1 in this region abolished its therapeutic effect, despite preserved activity in the rest of the brain, whereas inactivation in the right CA1 had no impact. Conversely, infusion of Alloswitch-1 restricted to the left CA1 was sufficient to fully restore short-term memory. These results suggest that, in the context of AD pathology, mGlu5 signaling in the left hippocampus plays a dominant role in short-term memory processing.

The divergence between our findings and previous optogenetic studies, where both hemispheres were required for short-term memory, likely reflects methodological differences. Optogenetic targets genetically defined neuronal populations and modulates synaptic transmission at the network level, which may broadly impact circuit excitability. In contrast, photopharmacology enables reversible, spatially confined modulation of endogenous receptor function, allowing for selective manipulation of specific pathways, such as mGlu5 signaling, under pathological conditions. This precision allows more nuanced dissection of pathway-specific contributions under pathological conditions. Moreover, while prior optogenetic studies were conducted in healthy animals, our approach targeted a disease context, revealing compensatory circuit dynamics and receptor-level contributions that may be masked under normal conditions.

Mechanistically, the selective vulnerability of the left hippocampus may be driven by mGlu5-dependent effects of Aβ. *Ex vivo* electrophysiology has shown that Aβ preferentially impairs plasticity at left CA3–CA1 synapses, enhancing LTD and impairing LTP in an mGlu5-dependent manner (*O’Riordan et al., 2018; Shipton et al., 2022*). Our findings extend this work by showing that left-sided mGlu5 inhibition normalizes downstream signaling, including Fyn, Pyk2, and GSK-3β, and reduces tau pathology. In J20 mice, Pyk2 and GSK-3β were activated in the left hippocampus, whereas Fyn activation was unexpectedly reduced at this stage, despite our previous report of elevated Fyn expression at 9 months (*Canet et al., 2025*). This reduction may reflect an early compensatory mechanism, as previously suggested in younger J20 mice through increased STEP phosphatase activity (*Chin et al., 2005*). Even though Fyn activation was reduced, left-sided Alloswitch-1 administration restored its activity to WT levels. Among the kinases studied, GSK-3β showed clear hemispheric asymmetry, with increased activation specifically in the left hippocampus, where we also observed a trend toward elevated levels of soluble Aβ_1–42_. Although asymmetry in rodent AD models remains underreported, similar lateralized patterns have been described: in TgF344-AD rats, increased microglial activity in the left hippocampus associates with spatial and memory deficits (*Sagalajev et al., 2023*), and multiple transgenic AD mice show asymmetric amyloid plaque deposition linked to lateralized neuroinflammation (*Sacher et al., 2020*). Together, these data suggest that hemispheric asymmetry is a reproducible feature in AD models, warranting greater consideration in disease research and therapeutic development.

Notably, despite this pronounced left-sided pathology and the unilateral nature of our intervention, we observed bilateral rescue effects on kinase signaling and tau pathology. This bilateral rescue likely reflects functional interhemispheric connectivity within CA1. Recent anatomical and functional studies have revealed previously unappreciated commissural projections between dorsal CA1 regions (*De León Reyes et al., 2024; Zhou et al., 2017*). Disruption of these connections impairs spatial memory, emphasizing their importance in coordinating hippocampal function. Such projections may mediate the spread of pathological signals and, conversely, enable therapeutic modulation across hemispheres.

Importantly, our work underscores the translational value of photopharmacology. Alloswitch-1 allowed us to isolate the role of mGlu5 with subregional precision, revealing the left CA1 as a critical therapeutic node. However, the compound’s basal activity in the dark limits its clinical applicability. To overcome this, we are developing next-generation photoactivatable ligands that remain inert until light-triggered, offering spatiotemporal precision with minimal off-target effects.

Together, our findings establish that mGlu5 function is lateralized in the hippocampus and that its left-sided disruption contributes to AD-related memory impairments. Spatially targeted inhibition in this region is sufficient to restore memory and mitigate pathology, even across hemispheres. These results highlight the therapeutic potential of precision neuromodulation strategies and point to the importance of considering functional lateralization in future AD interventions.

## Supporting information

SFig.2

SFig.3

## ACKNOWLEDGEMENT

Part of the figures were created with BioRender.com. The authors are thankful to Elisabeth HUETTER and the animal facility staff (CECEMA, University of Montpellier, France) for their daily assistance. Parts of this study were supported by INSERM, CNRS and University of Montpellier annual resources (France), and by grants from the “Fondation pour la Recherche Médicale" (n° MND202003011477-OPA, LG, CG); and the “University of Montpellier Program of Excellence” (n° MUSE-AAP20REC-FRS09-GAiA, LG, CG). LG was supported by a grant from the “Agence Nationale de la Recherche” (n° ANR-AAP2022-R22102FF-EpiNeurAge). CD was supported by a grant from the “Agence Nationale de la Recherche” under the program “Investissements d’Avenir” (ANR-11-LABEX-0021-LipSTIC). AL received funding from Ministerio de Ciencia e Innovación, Agencia Estatal de Investigación 10.13039/501100011033 and ERDF A way of making Europe (Project PID2020-120499RB-I00), and the Catalan government (2021 SGR 00508). The authors thank Fanny Malhaire (IGF), Lourdes Muñoz, Carme Serra and Juanlo Catena from SimChem (IQAC-CSIC) for technical support and Anaëlle Dumazer (IGF) for proofreading the final manuscript.

## AUTHORS CONTRIBUTIONS

LG and CZ designed the research; CZ, MV and MM performed the experiments; AL and XGS synthetized and provided Alloswitch-1 compound; CZ and LG analyzed data. All authors discussed the results. CZ and LG wrote the first draft of the manuscript, which was revised by all authors. LG, CG and AL acquired fundings. All authors read and approved the final manuscript.

## CONFLICT OF INTEREST

All authors declare that they have no competing interests.

---

**SUPPLEMENTAL FIGURE 1**

**A-B.** Asymmetrical levels of soluble Aβ_1-42_ and activated GSK-3β in the hippocampus of J20 mice. **A.** Soluble Aβ_1-42_ levels were measured in the **left** and **right** hippocampus by ELISA and expressed as the left/right ratio. A one-sample t-test was conducted to compare the sample mean to the theoretical value of 1.0. t(4) = 1.390 (ns). **B.** GSK-3β activation were measured in the left and right hippocampus by Western blot (Phospho-GSK-3β / total GSK-3β, Fig.4E-F) and expressed as the left/right ratio. A one-sample t-test was conducted to compare the sample mean to the theoretical value of 1.0. t(5) = 5.493 (p< 0.01). Orange dots represent individual animals.

**SUPPLEMENTAL FIGURE 2**

**A.** *Intraperitoneal (i.p.) injection protocol.* Working memory performance was assessed in a Y-maze 40 minutes after i.p. injection of Alloswitch-1 or vehicle. Mice were allowed to freely explore all three arms of the maze for 8 minutes. Spontaneous alternation behavior, which reflects short-term working memory, is driven by the innate tendency of rodents to explore novel environments. Alternation performance was calculated as: (number of spontaneous alternations / [total arm entries – 2]) × 100 and expressed as a percentage. **B.** Effect of *(i.p.)* injection of Alloswitch-1 (Allo) or vehicle (Veh) on working memory performance. One-way ANOVA followed by Dunnett’s post hoc test revealed a significant treatment effect (F_3,25_ = 5.43, *p* = 0.0051). * p< 0.05 *vs.* J20-Veh group. **C.** No effect of optrode implantation on working memory performance after *(i.p.)* injection of Alloswitch-1 (Allo, 20 mg/kg) in J20 mice. An unpaired two-tailed t-test revealed no significant difference between the two groups, t(14) = 0.058 (ns). All data are presented as box-and-whisker plots (min to max, with median), orange dots represent individual animals. Note that a part of this figure was created in BioRender.

**SUPPLEMENTAL FIGURE 3**

Effect of Alloswitch-1 infusion (300 nM) into the **left** (L) or **right** (R) hippocampus on mGlu5 expression and activation of Fyn kinase. For experimental protocol see Fig.3C. mGlu5 protein levels (132 kDa) in the **left** (**A,C**) and **right** (**B,D**) hippocampus were measured by Western blot and normalized to β-tubulin (β-tub, 50 kDa). Data are expressed as the mGlu5/β-tub ratio. One-way ANOVA followed by Dunnett’s multiple comparisons: **C**: F_3,23_ = 4.48 (p < 0.01); **D**: F_3,23_ = 0.36 (ns). * p< 0.05 *vs*. J20-Veh L group. Phospho-Fyn protein levels (59 kDa) in the **left** (**A,E**) and **right** (**B,F**) hippocampus, normalized to total Fyn levels (59 kDa). Data are expressed as the Phospho-Fyn/Fyn ratio. One-way ANOVA followed by Dunnett’s multiple comparisons: **E**: F_3,23_ = 13.23 (p < 0.0001); **F**: F_3,23_ = 7.99 (p < 0.001). * p< 0.05 and ** p< 0.01 *vs*. J20-Veh L group. All data are presented as box-and-whisker plots (min to max, with median), orange dots represent individual animals. Note that a part of this figure was created in BioRender.

**SUPPLEMENTAL FIGURE 4**

Western blot images for phosphorylated Pyk2 (p-Pyk2, 116 kDa) (**A**) and Pyk2 (116 kDa) (**B**) corresponding to the quantification of the Fig.4C-D. Lanes containing the molecular weight ladder are labeled (**L**), with the corresponding membrane number indicated below in red. Lanes corresponding to samples from the different experimental groups are labeled in black. **WL** and **WR** denote the left and right hippocampi of wild-type mice, respectively. For J20 mice, the first letter (**J**) denotes the genotype (J20). The second letter indicates the treatment: (**V**) for vehicle and (**A**) for Alloswitch-1. The third letter specifies the infused hippocampal side (**L** or **R**), and the fourth letter the dissected side (**L** or **R**).

**SUPPLEMENTAL FIGURE 5**

Western blot images for phosphorylated GSK-3β (p-GSK-3β, 46 kDa) (**A**) and GSK-3β (46 kDa) (**B**) corresponding to the quantification of the Fig.4E-F. Lanes containing the molecular weight ladder are labeled (**L**), with the corresponding membrane number indicated below in red. Lanes corresponding to samples from the different experimental groups are labeled in black. **WL** and **WR** denote the left and right hippocampi of wild-type mice, respectively. For J20 mice, the first letter (**J**) denotes the genotype (J20). The second letter indicates the treatment: (**V**) for vehicle and (**A**) for Alloswitch-1. The third letter specifies the infused hippocampal side (**L** or **R**), and the fourth letter the dissected side (**L** or **R**).

**SUPPLEMENTAL FIGURE 6**

Western blot images for phosphorylated Tau (p-Tau, AT180, 60 kDa) (**A**) and total Tau (t-Tau, 60 kDa) (**B**) corresponding to the quantification of the Fig.4G-H. Lanes containing the molecular weight ladder are labeled (**L**), with the corresponding membrane number indicated below in red. Lanes corresponding to samples from the different experimental groups are labeled in black. **WL** and **WR** denote the left and right hippocampi of wild-type mice, respectively. For J20 mice, the first letter (**J**) denotes the genotype (J20). The second letter indicates the treatment: (**V**) for vehicle and (**A**) for Alloswitch-1. The third letter specifies the infused hippocampal side (**L** or **R**), and the fourth letter the dissected side (**L** or **R**).

**SUPPLEMENTAL FIGURE 7**

Western blot images for mGlu5 (132 kDa) (**A**) and β-tubulin (β-tub, 50 kDa) (**B**) corresponding to the quantification of the supplemental Fig.3C-D. Lanes containing the molecular weight ladder are labeled (**L**), with the corresponding membrane number indicated below in red. Lanes corresponding to samples from the different experimental groups are labeled in black. **WL** and **WR** denote the left and right hippocampi of wild-type mice, respectively. For J20 mice, the first letter (**J**) denotes the genotype (J20). The second letter indicates the treatment: (**V**) for vehicle and (**A**) for Alloswitch-1. The third letter specifies the infused hippocampal side (**L** or **R**), and the fourth letter the dissected side (**L** or **R**).

**SUPPLEMENTAL FIGURE 8**

Western blot images for Phospho-Fyn (p-Fyn, 59 kDa) (**A**) and Fyn (59 kDa) (**B**) corresponding to the quantification of the supplemental Fig.3E-F. Lanes containing the molecular weight ladder are labeled (**L**), with the corresponding membrane number indicated below in red. Lanes corresponding to samples from the different experimental groups are labeled in black. **WL** and **WR** denote the left and right hippocampi of wild-type mice, respectively. For J20 mice, the first letter (**J**) denotes the genotype (J20). The second letter indicates the treatment: (**V**) for vehicle and (**A**) for Alloswitch-1. The third letter specifies the infused hippocampal side (**L** or **R**), and the fourth letter the dissected side (**L** or **R**).

## Notes

### Competing Interest Statement

The authors have declared no competing interest.

## REFERENCES

Abd-Elrahman, K.S., Ferguson, S.S.G., 2022. Noncanonical Metabotropic Glutamate Receptor 5 Signaling in Alzheimer’s Disease. Annu. Rev. Pharmacol. Toxicol. 62, 235–254. 10.1146/annurev-pharmtox-021821-091747

Bikbaev, A., Manahan-Vaughan, D., 2017. Metabotropic glutamate receptor, mGlu5, regulates hippocampal synaptic plasticity and is required for tetanisation-triggered changes in theta and gamma oscillations. Neuropharmacology 115, 20–29. 10.1016/j.neuropharm.2016.06.004

Brody, A.H., Strittmatter, S.M., 2018. Chapter Thirteen - Synaptotoxic Signaling by Amyloid Beta Oligomers in Alzheimer’s Disease Through Prion Protein and mGluR5, in: Pasternak, G.W., Coyle, J.T. (Eds.), Advances in Pharmacology, Apprentices to Genius: A Tribute to Solomon H. Snyder. Academic Press, pp. 293–323. 10.1016/bs.apha.2017.09.007

Budgett, R.F., Bakker, G., Sergeev, E., Bennett, K.A., Bradley, S.J., 2022. Targeting the Type 5 Metabotropic Glutamate Receptor: A Potential Therapeutic Strategy for Neurodegenerative Diseases? Frontiers in Pharmacology 13.

Bukke, V.N., Archana, M., Villani, R., Romano, A.D., Wawrzyniak, A., Balawender, K., Orkisz, S., Beggiato, S., Serviddio, G., Cassano, T., 2020. The Dual Role of Glutamatergic Neurotransmission in Alzheimer’s Disease: From Pathophysiology to Pharmacotherapy. IJMS 21, 7452. 10.3390/ijms21207452

Canet, G., Zussy, C., Vitalis, M., Morin, F., Chevallier, N., Hunt, H., Claeysen, S., Blaquière, M., Marchi, N., Planel, E., Meijer, O.C., Desrumaux, C., Givalois, L., 2025. Advancing Alzheimer’s disease pharmacotherapy: efficacy of glucocorticoid modulation with dazucorilant (CORT113176) in preclinical mouse models. British J Pharmacology 182, 1930–1956. 10.1111/bph.17457

Chin, J., Palop, J.J., Puoliväli, J., Massaro, C., Bien-Ly, N., Gerstein, H., Scearce-Levie, K., Masliah, E., Mucke, L., 2005. Fyn Kinase Induces Synaptic and Cognitive Impairments in a Transgenic Mouse Model of Alzheimer’s Disease. J. Neurosci. 25, 9694–9703. 10.1523/JNEUROSCI.2980-05.2005

Crestini, A., Santilli, F., Martellucci, S., Carbone, E., Sorice, M., Piscopo, P., Mattei, V., 2022. Prions and Neurodegenerative Diseases: A Focus on Alzheimer’s Disease. JAD 85, 503–518. 10.3233/JAD-215171

De Bundel, D., Zussy, C., Espallergues, J., Gerfen, C.R., Girault, J.-A., Valjent, E., 2016. Dopamine D2 receptors gate generalization of conditioned threat responses through mTORC1 signaling in the extended amygdala. Mol Psychiatry 21, 1545–1553. 10.1038/mp.2015.210

De León Reyes, N.S., Bortolozzo-Gleich, M.H., Nomura, Y., García Fregola, C., Nieto, M., Gogos, J.A., Leroy, F., 2024. Interhemispheric CA1 projections support spatial cognition and are affected in a mouse model of the 22q11.2 deletion syndrome. 10.1101/2024.09.05.611389

Derflinger, S., Sorg, C., Gaser, C., Myers, N., Arsic, M., Kurz, A., Zimmer, C., Wohlschläger, A., Mühlau, M., 2011. Grey-Matter Atrophy in Alzheimer’s Disease is Asymmetric but not Lateralized. JAD 25, 347–357. 10.3233/JAD-2011-110041

DeTure, M.A., Dickson, D.W., 2019. The neuropathological diagnosis of Alzheimer’s disease. Mol Neurodegeneration 14, 32. 10.1186/s13024-019-0333-5

Dohler, F., Sepulveda-Falla, D., Krasemann, S., Altmeppen, H., Schlüter, H., Hildebrand, D., Zerr, I., Matschke, J., Glatzel, M., 2014. High molecular mass assemblies of amyloid-β oligomers bind prion protein in patients with Alzheimer’s disease. Brain 137, 873–886. 10.1093/brain/awt375

Dourlen, P., Fernandez-Gomez, F.J., Dupont, C., Grenier-Boley, B., Bellenguez, C., Obriot, H., Caillierez, R., Sottejeau, Y., Chapuis, J., Bretteville, A., Abdelfettah, F., Delay, C., Malmanche, N., Soininen, H., Hiltunen, M., Galas, M.-C., Amouyel, P., Sergeant, N., Buée, L., Lambert, J.-C., Dermaut, B., 2017. Functional screening of Alzheimer risk loci identifies PTK2B as an in vivo modulator and early marker of Tau pathology. Mol Psychiatry 22, 874–883. 10.1038/mp.2016.59

Fang, X.T., Eriksson, J., Antoni, G., Yngve, U., Cato, L., Lannfelt, L., Sehlin, D., Syvänen, S., 2017. Brain mGluR5 in mice with amyloid beta pathology studied with in vivo [11C]ABP688 PET imaging and ex vivo immunoblotting. Neuropharmacology 113, 293–300. 10.1016/j.neuropharm.2016.10.009

Fox, N.C., Warrington, E.K., Freeborough, P.A., Hartikainen, P., Kennedy, A.M., Stevens, J.M., Rossor, M.N., 1996. Presymptomatic hippocampal atrophy in Alzheimer’s disease. A longitudinal MRI study. Brain 119 ( Pt 6), 2001–2007. 10.1093/brain/119.6.2001

Frings, L., Hellwig, S., Spehl, T.S., Bormann, T., Buchert, R., Vach, W., Minkova, L., Heimbach, B., Klöppel, S., Meyer, P.T., 2015. Asymmetries of amyloid-β burden and neuronal dysfunction are positively correlated in Alzheimer’s disease. Brain 138, 3089–3099. 10.1093/brain/awv229

Gómez-Santacana, X., Panarello, S., Rovira, X., Llebaria, A., 2022. Photoswitchable allosteric modulators for metabotropic glutamate receptors. Current Opinion in Pharmacology 66, 102266. 10.1016/j.coph.2022.102266

Gómez-Santacana, X., Pittolo, S., Rovira, X., Lopez, M., Zussy, C., Dalton, J.A.R., Faucherre, A., Jopling, C., Pin, J.-P., Ciruela, F., Goudet, C., Giraldo, J., Gorostiza, P., Llebaria, A., 2017. Illuminating Phenylazopyridines To Photoswitch Metabotropic Glutamate Receptors: From the Flask to the Animals. ACS Cent. Sci. 3, 81–91. 10.1021/acscentsci.6b00353

Haas, L.T., Salazar, S.V., Kostylev, M.A., Um, J.W., Kaufman, A.C., Strittmatter, S.M., 2016. Metabotropic glutamate receptor 5 couples cellular prion protein to intracellular signalling in Alzheimer’s disease. Brain 139, 526–546. 10.1093/brain/awv356

Hagena, H., Manahan-Vaughan, D., 2022. Role of mGlu5 in Persistent Forms of Hippocampal Synaptic Plasticity and the Encoding of Spatial Experience. Cells 11, 3352. 10.3390/cells11213352

Hartigan, J.A., Xiong, W.C., Johnson, G.V., 2001. Glycogen synthase kinase 3beta is tyrosine phosphorylated by PYK2. Biochem Biophys Res Commun 284, 485–489. 10.1006/bbrc.2001.4986

Holcomb, L., Gordon, M.N., McGowan, E., Yu, X., Benkovic, S., Jantzen, P., Wright, K., Saad, I., Mueller, R., Morgan, D., Sanders, S., Zehr, C., O’Campo, K., Hardy, J., Prada, C.M., Eckman, C., Younkin, S., Hsiao, K., Duff, K., 1998. Accelerated Alzheimer-type phenotype in transgenic mice carrying both mutant amyloid precursor protein and presenilin 1 transgenes. Nat Med 4, 97–100. 10.1038/nm0198-097

Hu, N.-W., Nicoll, A.J., Zhang, D., Mably, A.J., O’Malley, T., Purro, S.A., Terry, C., Collinge, J., Walsh, D.M., Rowan, M.J., 2014. mGlu5 receptors and cellular prion protein mediate amyloid-β-facilitated synaptic long-term depression in vivo. Nat Commun 5, 3374. 10.1038/ncomms4374

Hughes, R.N., 2004. The value of spontaneous alternation behavior (SAB) as a test of retention in pharmacological investigations of memory. Neuroscience & Biobehavioral Reviews 28, 497–505. 10.1016/j.neubiorev.2004.06.006

Iaccarino, H.F., Singer, A.C., Martorell, A.J., Rudenko, A., Gao, F., Gillingham, T.Z., Mathys, H., Seo, J., Kritskiy, O., Abdurrob, F., Adaikkan, C., Canter, R.G., Rueda, R., Brown, E.N., Boyden, E.S., Tsai, L.-H., 2016. Gamma frequency entrainment attenuates amyloid load and modifies microglia. Nature 540, 230–235. 10.1038/nature20587

Kawakami, R., Shinohara, Y., Kato, Y., Sugiyama, H., Shigemoto, R., Ito, I., 2003. Asymmetrical Allocation of NMDA Receptor 2 Subunits in Hippocampal Circuitry. Science 300, 990–4. 10.1126/science.1082609.

Kitanishi, T., Ito, H.T., Hayashi, Y., Shinohara, Y., Mizuseki, K., Hikida, T., 2017. Network mechanisms of hippocampal laterality, place coding, and goal-directed navigation. The Journal of Physiological Sciences 67, 247–258. 10.1007/s12576-016-0502-z

Kohl, M.M., Shipton, O.A., Deacon, R.M., Rawlins, J.N.P., Deisseroth, K., Paulsen, O., 2011. Hemisphere-specific optogenetic stimulation reveals left-right asymmetry of hippocampal plasticity. Nat Neurosci 14, 1413–1415. 10.1038/nn.2915

Kolber, B.J., Montana, M.C., Carrasquillo, Y., Xu, J., Heinemann, S.F., Muglia, L.J., Gereau, R.W., 2010. Activation of Metabotropic Glutamate Receptor 5 in the Amygdala Modulates Pain-Like Behavior. Journal of Neuroscience 30, 8203–8213. 10.1523/JNEUROSCI.1216-10.2010

Kumar, A., Dhull, D.K., Mishra, P.S., 2015. Therapeutic potential of mGluR5 targeting in Alzheimer’s disease. Front. Neurosci. 9. 10.3389/fnins.2015.00215

Lambert, J.-C., Ibrahim-Verbaas, C.A., Harold, D., Naj, A.C., Sims, R., Jun, G., DeStefano, A.L., Bis, J.C., Beecham, G.W., Russo, G., Thornton-Wells, T.A., Jones, N., Smith, A.V., Chouraki, V., Thomas, C., Ikram, M.A., Zelenika, D., Vardarajan, B.N., Kamatani, Y., Lin, C.-F., Gerrish, A., Schmidt, H., Kunkle, B., Dunstan, M.L., Ruiz, A., Bihoreau, M.-T., Choi, S.-H., Reitz, C., Pasquier, F., Hollingworth, P., Ramirez, A., Hanon, O., Fitzpatrick, A.L., Buxbaum, J.D., Campion, D., Crane, P.K., Baldwin, C., Becker, T., Gudnason, V., Cruchaga, C., Craig, D., Amin, N., Berr, C., Lopez, O.L., De Jager, P.L., Deramecourt, V., Johnston, J.A., Evans, D., Lovestone, S., Letenneur, L., Morón, F.J., Rubinsztein, D.C., Eiriksdottir, G., Sleegers, K., Goate, A.M., Fiévet, N., Huentelman, M.J., Gill, M., Brown, K., Kamboh, M.I., Keller, L., Barberger-Gateau, P., McGuinness, B., Larson, E.B., Green, R., Myers, A.J., Dufouil, C., Todd, S., Wallon, D., Love, S., Rogaeva, E., Gallacher, J., St George-Hyslop, P., Clarimon, J., Lleo, A., Bayer, A., Tsuang, D.W., Yu, L., Tsolaki, M., Bossù, P., Spalletta, G., Proitsi, P., Collinge, J., Sorbi, S., Sanchez-Garcia, F., Fox, N.C., Hardy, J., Naranjo, M.C.D., Bosco, P., Clarke, R., Brayne, C., Galimberti, D., Mancuso, M., Matthews, F., Moebus, S., Mecocci, P., Del Zompo, M., Maier, W., Hampel, H., Pilotto, A., Bullido, M., Panza, F., Caffarra, P., Nacmias, B., Gilbert, J.R., Mayhaus, M., Lannfelt, L., Hakonarson, H., Pichler, S., Carrasquillo, M.M., Ingelsson, M., Beekly, D., Alvarez, V., Zou, F., Valladares, O., Younkin, S.G., Coto, E., Hamilton-Nelson, K.L., Gu, W., Razquin, C., Pastor, P., Mateo, I., Owen, M.J., Faber, K.M., Jonsson, P.V., Combarros, O., O’Donovan, M.C., Cantwell, L.B., Soininen, H., Blacker, D., Mead, S., Mosley, T.H., Bennett, D.A., Harris, T.B., Fratiglioni, L., Holmes, C., De Bruijn, R.F.A.G., Passmore, P., Montine, T.J., Bettens, K., Rotter, J.I., Brice, A., Morgan, K., Foroud, T.M., Kukull, W.A., Hannequin, D., Powell, J.F., Nalls, M.A., Ritchie, K., Lunetta, K.L., Kauwe, J.S.K., Boerwinkle, E., Riemenschneider, M., Boada, M., Hiltunen, M., Martin, E.R., Schmidt, R., Rujescu, D., Wang, L.-S., Dartigues, J.-F., Mayeux, R., Tzourio, C., Hofman, A., Nöthen, M.M., Graff, C., Psaty, B.M., Jones, L., Haines, J.L., Holmans, P.A., Lathrop, M., Pericak-Vance, M.A., Launer, L.J., Farrer, L.A., Van Duijn, C.M., Van Broeckhoven, C., Moskvina, V., Seshadri, S., Williams, J., Schellenberg, G.D., Amouyel, P., 2013. Meta-analysis of 74,046 individuals identifies 11 new susceptibility loci for Alzheimer’s disease. Nat Genet 45, 1452–1458. 10.1038/ng.2802

Laurén, J., Gimbel, D.A., Nygaard, H.B., Gilbert, J.W., Strittmatter, S.M., 2009. Cellular prion protein mediates impairment of synaptic plasticity by amyloid-β oligomers. Nature 457, 1128–1132. 10.1038/nature07761

Lee, M., Lee, H.-J., Park, I.S., Park, J.-A., Kwon, Y.J., Ryu, Y.H., Kim, C.H., Kang, J.H., Hyun, I.Y., Lee, K.C., Choi, J.Y., 2018. Aβ pathology downregulates brain mGluR5 density in a mouse model of Alzheimer. Neuropharmacology 133, 512–517. 10.1016/j.neuropharm.2018.02.003

Li, C., Götz, J., 2018. Pyk2 is a Novel Tau Tyrosine Kinase that is Regulated by the Tyrosine Kinase Fyn. JAD 64, 205–221. 10.3233/JAD-180054

Lue, L.F., Kuo, Y.M., Roher, A.E., Brachova, L., Shen, Y., Sue, L., Beach, T., Kurth, J.H., Rydel, R.E., Rogers, J., 1999. Soluble amyloid beta peptide concentration as a predictor of synaptic change in Alzheimer’s disease. Am J Pathol 155, 853–862. 10.1016/s0002-9440(10)65184-x

Mansuy, M., Baille, S., Canet, G., Borie, A., Vignes, M., Perrier, V., Chevallier, N., Guern, L., Deckert, V., Lagrost, L., Givalois, L., Desrumaux, C., 2018. Deletion of plasma Phospholipid Transfer Protein (PLTP) increases microglial phagocytosis and reduces cerebral amyloid-β deposition in the J20 mouse model of Alzheimer’s disease. Oncotarget Vol. 9, 19688–19703.

McLean, C.A., Cherny, R.A., Fraser, F.W., Fuller, S.J., Smith, M.J., Vbeyreuther, K., Bush, A.I., Masters, C.L., 1999. Soluble pool of Aβ amyloid as a determinant of severity of neurodegeneration in Alzheimer’s disease. Annals of Neurology 46, 860–866. 10.1002/1531-8249(199912)46:6<860::AID-ANA8>3.0.CO;2-M

Mecca, A.P., McDonald, J.W., Michalak, H.R., Godek, T.A., Harris, J.E., Pugh, E.A., Kemp, E.C., Chen, M.-K., Salardini, A., Nabulsi, N.B., Lim, K., Huang, Y., Carson, R.E., Strittmatter, S.M., Van Dyck, C.H., 2020. PET imaging of mGluR5 in Alzheimer’s disease. Alz Res Therapy 12, 15. 10.1186/s13195-020-0582-0

Miller, J., Watrous, A.J., Tsitsiklis, M., Lee, S.A., Sheth, S.A., Schevon, C.A., Smith, E.H., Sperling, M.R., Sharan, A., Asadi-Pooya, A.A., Worrell, G.A., Meisenhelter, S., Inman, C.S., Davis, K.A., Lega, B., Wanda, P.A., Das, S.R., Stein, J.M., Gorniak, R., Jacobs, J., 2018. Lateralized hippocampal oscillations underlie distinct aspects of human spatial memory and navigation. Nat Commun 9, 2423. 10.1038/s41467-018-04847-9

Miller, S.L., Fenstermacher, E., Bates, J., Blacker, D., Sperling, R.A., Dickerson, B.C., 2007. Hippocampal activation in adults with mild cognitive impairment predicts subsequent cognitive decline. Journal of Neurology, Neurosurgery & Psychiatry 79, 630–635. 10.1136/jnnp.2007.124149

Mucke, L., Masliah, E., Yu, G.-Q., Mallory, M., Rockenstein, E.M., Tatsuno, G., Hu, K., Kholodenko, D., Johnson-Wood, K., McConlogue, L., 2000. High-Level Neuronal Expression of Aβ1–42 in Wild-Type Human Amyloid Protein Precursor Transgenic Mice: Synaptotoxicity without Plaque Formation. J. Neurosci. 20, 4050–4058. 10.1523/JNEUROSCI.20-11-04050.2000

Mukherjee, S., Manahan-Vaughan, D., 2013. Role of metabotropic glutamate receptors in persistent forms of hippocampal plasticity and learning. Neuropharmacology 66, 65–81. 10.1016/j.neuropharm.2012.06.005

Müller Herde, A., Schibli, R., Weber, M., Ametamey, S.M., 2019. Metabotropic glutamate receptor subtype 5 is altered in LPS-induced murine neuroinflammation model and in the brains of AD and ALS patients. Eur J Nucl Med Mol Imaging 46, 407–420. 10.1007/s00259-018-4179-9

Ng, A.N., Salter, E.W., Georgiou, J., Bortolotto, Z.A., Collingridge, G.L., 2023. Amyloid-β1-42 oligomers enhance mGlu5R-dependent synaptic weakening via NMDAR activation and complement C5aR1 signaling. iScience 26, 108412. 10.1016/j.isci.2023.108412

Notartomaso, S., Antenucci, N., Mazzitelli, M., Rovira, X., Boccella, S., Ricciardi, F., Liberatore, F., Gomez-Santacana, X., Imbriglio, T., Cannella, M., Zussy, C., Luongo, L., Maione, S., Goudet, C., Battaglia, G., Llebaria, A., Nicoletti, F., Neugebauer, V., 2024. A “double-edged” role for type-5 metabotropic glutamate receptors in pain disclosed by light-sensitive drugs. Elife 13, e94931. 10.7554/eLife.94931

Oddo, S., Caccamo, A., Tran, L., Lambert, M.P., Glabe, C.G., Klein, W.L., LaFerla, F.M., 2006. Temporal profile of amyloid-beta (Abeta) oligomerization in an in vivo model of Alzheimer disease. A link between Abeta and tau pathology. J Biol Chem 281, 1599–1604. 10.1074/jbc.M507892200

O’Riordan, K.J., Hu, N.-W., Rowan, M.J., 2018. Aß Facilitates LTD at Schaffer Collateral Synapses Preferentially in the Left Hippocampus. Cell Reports 22, 2053–2065. 10.1016/j.celrep.2018.01.085

Panarello, S., Rovira, X., Llebaria, A., Gómez-Santacana, X., 2022. Photopharmacology of G -Protein-Coupled Receptors, in: Molecular Photoswitches. John Wiley & Sons, Ltd, pp. 921–944. 10.1002/9783527827626.ch37

Pittolo, S., Gómez-Santacana, X., Eckelt, K., Rovira, X., Dalton, J., Goudet, C., Pin, J.-P., Llobet, A., Giraldo, J., Llebaria, A., Gorostiza, P., 2014. An allosteric modulator to control endogenous G protein-coupled receptors with light. Nat Chem Biol 10, 813–815. 10.1038/nchembio.1612

Privitera, L., Hogg, E.L., Lopes, M., Domingos, L.B., Gaestel, M., Müller, J., Wall, M.J., Corrêa, S.A.L., 2022. The MK2 cascade mediates transient alteration in mGluR-LTD and spatial learning in a murine model of Alzheimer’s disease. Aging Cell 21, e13717. 10.1111/acel.13717

Rajani, V., Sengar, A.S., Salter, M.W., 2021. Src and Fyn regulation of NMDA receptors in health and disease. Neuropharmacology 193, 108615. 10.1016/j.neuropharm.2021.108615

Renner, M., Lacor, P.N., Velasco, P.T., Xu, J., Contractor, A., Klein, W.L., Triller, A., 2010. Deleterious Effects of Amyloid β Oligomers Acting as an Extracellular Scaffold for mGluR5. Neuron 66, 739–754. 10.1016/j.neuron.2010.04.029

Ricart-Ortega, M., Berizzi, A.E., Pereira, V., Malhaire, F., Catena, J., Font, J., Gómez-Santacana, X., Muñoz, L., Zussy, C., Serra, C., Rovira, X., Goudet, C., Llebaria, A., 2020. Mechanistic Insights into Light-Driven Allosteric Control of GPCR Biological Activity. ACS Pharmacol. Transl. Sci. 3, 883–895. 10.1021/acsptsci.0c00054

Roberson, E.D., Halabisky, B., Yoo, J.W., Yao, J., Chin, J., Yan, F., Wu, T., Hamto, P., Devidze, N., Yu, G.-Q., Palop, J.J., Noebels, J.L., Mucke, L., 2011. Amyloid-β/Fyn–Induced Synaptic, Network, and Cognitive Impairments Depend on Tau Levels in Multiple Mouse Models of Alzheimer’s Disease. J. Neurosci. 31, 700– 711. 10.1523/JNEUROSCI.4152-10.2011

Sacher, C., Blume, T., Beyer, L., Biechele, G., Sauerbeck, J., Eckenweber, F., Deussing, M., Focke, C., Parhizkar, S., Lindner, S., Gildehaus, F.-J., Von Ungern-Sternberg, B., Baumann, K., Tahirovic, S., Kleinberger, G., Willem, M., Haass, C., Bartenstein, P., Cumming, P., Rominger, A., Herms, J., Brendel, M., 2020. Asymmetry of Fibrillar Plaque Burden in Amyloid Mouse Models. J Nucl Med 61, 1825–1831. 10.2967/jnumed.120.242750

Sagalajev, B., Lennartz, L., Vieth, L., Gunawan, C.T., Neumaier, B., Drzezga, A., Visser-Vandewalle, V., Endepols, H., Sesia, T., 2023. TgF344-AD Rat Model of Alzheimer’s Disease: Spatial Disorientation and Asymmetry in Hemispheric Neurodegeneration. Journal of Alzheimer’s Disease Reports 7, 1085–1094. 10.3233/adr-230038

Salardini, E., O’Dell, R.S., Tchorz, E., Nabulsi, N.B., Huang, Y., Carson, R.E., van Dyck, C.H., Mecca, A.P., 2024. Assessment of the relationship between synaptic density and metabotropic glutamate receptors in early Alzheimer’s disease: a multi-tracer PET study. bioRxiv 2024.09.21.614277. 10.1101/2024.09.21.614277

Scahill, R.I., Schott, J.M., Stevens, J.M., Rossor, M.N., Fox, N.C., 2002. Mapping the evolution of regional atrophy in Alzheimer’s disease: unbiased analysis of fluid-registered serial MRI. Proc Natl Acad Sci U S A 99, 4703–4707. 10.1073/pnas.052587399

Selkoe, D.J., 2001. Alzheimer’s Disease: Genes, Proteins, and Therapy. Physiological Reviews 81, 741–766. 10.1152/physrev.2001.81.2.741

Sengupta, A., Kabat, J., Novak, M., Wu, Q., Grundke-Iqbal, I., Iqbal, K., 1998. Phosphorylation of Tau at Both Thr 231 and Ser 262 Is Required for Maximal Inhibition of Its Binding to Microtubules. Archives of Biochemistry and Biophysics 357, 299–309. 10.1006/abbi.1998.0813

Shankar, G.M., Li, S., Mehta, T.H., Garcia-Munoz, A., Shepardson, N.E., Smith, I., Brett, F.M., Farrell, M.A., Rowan, M.J., Lemere, C.A., Regan, C.M., Walsh, D.M., Sabatini, B.L., Selkoe, D.J., 2008. Amyloid-beta protein dimers isolated directly from Alzheimer’s brains impair synaptic plasticity and memory. Nat Med 14, 837–842. 10.1038/nm1782

Shinohara, Y., Hirase, H., 2009. Size and receptor density of glutamatergic synapses: a viewpoint from left-right asymmetry of CA3-CA1 connections. Frontiers in Neuroanatomy 3.

Shinohara, Y., Hirase, H., Watanabe, M., Itakura, M., Takahashi, M., Shigemoto, R., 2008. Left-right asymmetry of the hippocampal synapses with differential subunit allocation of glutamate receptors. Proc. Natl. Acad. Sci. U.S.A. 105, 19498–19503. 10.1073/pnas.0807461105

Shipton, O.A., El-Gaby, M., Apergis-Schoute, J., Deisseroth, K., Bannerman, D.M., Paulsen, O., Kohl, M.M., 2014. Left–right dissociation of hippocampal memory processes in mice. Proc. Natl. Acad. Sci. U.S.A. 111, 15238–15243. 10.1073/pnas.1405648111

Shipton, O.A., Tang, C.S., Paulsen, O., Vargas-Caballero, M., 2022. Differential vulnerability of hippocampal CA3-CA1 synapses to Aβ. Acta Neuropathologica Communications 10, 45. 10.1186/s40478-022-01350-7

Spurrier, J., Nicholson, L., Fang, X.T., Stoner, A.J., Toyonaga, T., Holden, D., Siegert, T.R., Laird, W., Allnutt, M.A., Chiasseu, M., Brody, A.H., Takahashi, H., Nies, S.H., Pérez-Cañamás, A., Sadasivam, P., Lee, S., Li, S., Zhang, L., Huang, Y.H., Carson, R.E., Cai, Z., Strittmatter, S.M., 2022. Reversal of synapse loss in Alzheimer mouse models by targeting mGluR5 to prevent synaptic tagging by C1Q. Science Translational Medicine 14, eabi8593. 10.1126/scitranslmed.abi8593

Treyer, V., Gietl, A.F., Suliman, H., Gruber, E., Meyer, R., Buchmann, A., Johayem, A., Unschuld, P.G., Nitsch, R.M., Buck, A., Ametamey, S.M., Hock, C., 2020. Reduced uptake of [11C]-ABP688, a PET tracer for metabolic glutamate receptor 5 in hippocampus and amygdala in Alzheimer’s dementia. Brain and Behavior 10, e01632. 10.1002/brb3.1632

Tyrer, A., Gilbert, J.R., Adams, S., Stiles, A.B., Bankole, A.O., Gilchrist, I.D., Moran, R.J., 2020. Lateralized memory circuit dropout in Alzheimer’s disease patients. Brain Communications 2, fcaa212. 10.1093/braincomms/fcaa212

Um, J.W., Kaufman, A.C., Kostylev, M., Heiss, J.K., Stagi, M., Takahashi, H., Kerrisk, M.E., Vortmeyer, A., Wisniewski, T., Koleske, A.J., Gunther, E.C., Nygaard, H.B., Strittmatter, S.M., 2013. Metabotropic Glutamate Receptor 5 Is a Coreceptor for Alzheimer Aβ Oligomer Bound to Cellular Prion Protein. Neuron 79, 887–902. 10.1016/j.neuron.2013.06.036

Um, J.W., Nygaard, H.B., Heiss, J.K., Kostylev, M.A., Stagi, M., Vortmeyer, A., Wisniewski, T., Gunther, E.C., Strittmatter, S.M., 2012. Alzheimer amyloid-β oligomer bound to postsynaptic prion protein activates Fyn to impair neurons. Nat Neurosci 15, 1227–1235. 10.1038/nn.3178

Valdivia, G., Ardiles, A.O., Idowu, A., Salazar, C., Lee, H.-K., Gallagher, M., Palacios, A.G., Kirkwood, A., 2023. mGluR-dependent plasticity in rodent models of Alzheimer’s disease. Front Synaptic Neurosci 15, 1123294. 10.3389/fnsyn.2023.1123294

Velema, W.A., Szymanski, W., Feringa, B.L., 2014. Photopharmacology: Beyond Proof of Principle. J. Am. Chem. Soc. 136, 2178–2191. 10.1021/ja413063e

Weise, C.M., Chen, K., Chen, Y., Kuang, X., Savage, C.R., Reiman, E.M., 2018. Left lateralized cerebral glucose metabolism declines in amyloid-β positive persons with mild cognitive impairment. NeuroImage: Clinical 20, 286–296. 10.1016/j.nicl.2018.07.016

You, H., Tsutsui, S., Hameed, S., Kannanayakal, T.J., Chen, L., Xia, P., Engbers, J.D.T., Lipton, S.A., Stys, P.K., Zamponi, G.W., 2012. Aβ neurotoxicity depends on interactions between copper ions, prion protein, and N -methyl-D -aspartate receptors. Proc. Natl. Acad. Sci. U.S.A. 109, 1737–1742. 10.1073/pnas.1110789109

Yu, X., Zhang, Y., Cai, Y., Rong, N., Li, R., Shi, R., Wei, M., Jiang, J., Han, Y., 2024a. Asymmetrical patterns of β-amyloid deposition and cognitive changes in Alzheimer’s disease: the SILCODE study. Cerebral Cortex 34. 10.1093/cercor/bhae485

Yu, X., Zhang, Y., Cai, Y., Rong, N., Li, R., Shi, R., Wei, M., Jiang, J., Han, Y., 2024b. Asymmetrical patterns of β-amyloid deposition and cognitive changes in Alzheimer’s disease: the SILCODE study. Cerebral Cortex 34, bhae485. 10.1093/cercor/bhae485

Zhou, H., Xiong, G.-J., Jing, L., Song, N.-N., Pu, D.-L., Tang, X., He, X.-B., Xu, F.-Q., Huang, J.-F., Li, L.-J., Richter-Levin, G., Mao, R.-R., Zhou, Q.-X., Ding, Y.-Q., Xu, L., 2017. The interhemispheric CA1 circuit governs rapid generalisation but not fear memory. Nat Commun 8, 2190. 10.1038/s41467-017-02315-4

Zussy, C., Gómez-Santacana, X., Rovira, X., De Bundel, D., Ferrazzo, S., Bosch, D., Asede, D., Malhaire, F., Acher, F., Giraldo, J., Valjent, E., Ehrlich, I., Ferraguti, F., Pin, J.-P., Llebaria, A., Goudet, C., 2018. Dynamic modulation of inflammatory pain-related affective and sensory symptoms by optical control of amygdala metabotropic glutamate receptor 4. Mol Psychiatry 23, 509–520. 10.1038/mp.2016.223

Zussy, C., John, R., Urgin, T., Otaegui, L., Vigor, C., Acar, N., Canet, G., Vitalis, M., Morin, F., Planel, E., Oger, C., Durand, T., Rajshree, S.L., Givalois, L., Devarajan, P.V., Desrumaux, C., 2022. Intranasal Administration of Nanovectorized Docosahexaenoic Acid (DHA) Improves Cognitive Function in Two Complementary Mouse Models of Alzheimer’s Disease. Antioxidants 11, 838. 10.3390/antiox11050838

