## Supplementary figures and images for "Shedding light on left hippocampal mGlu5 in Alzheimer’s disease"

### SFig.2

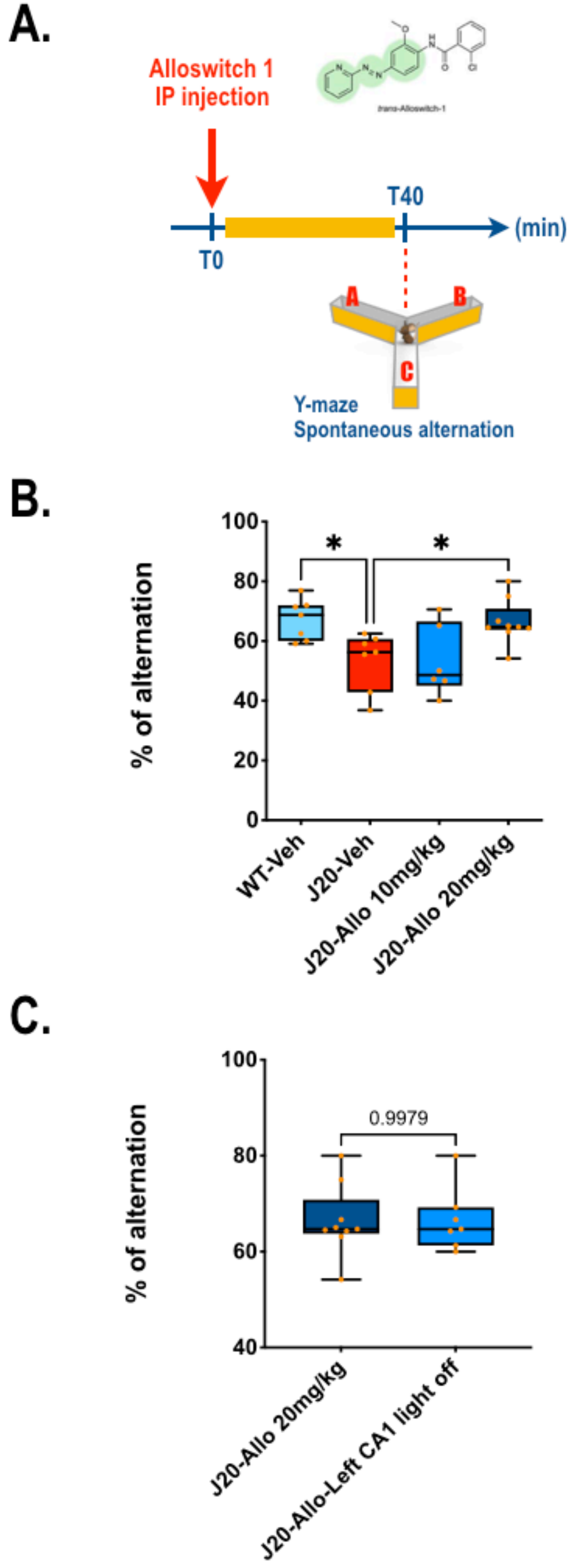

### SFig.3

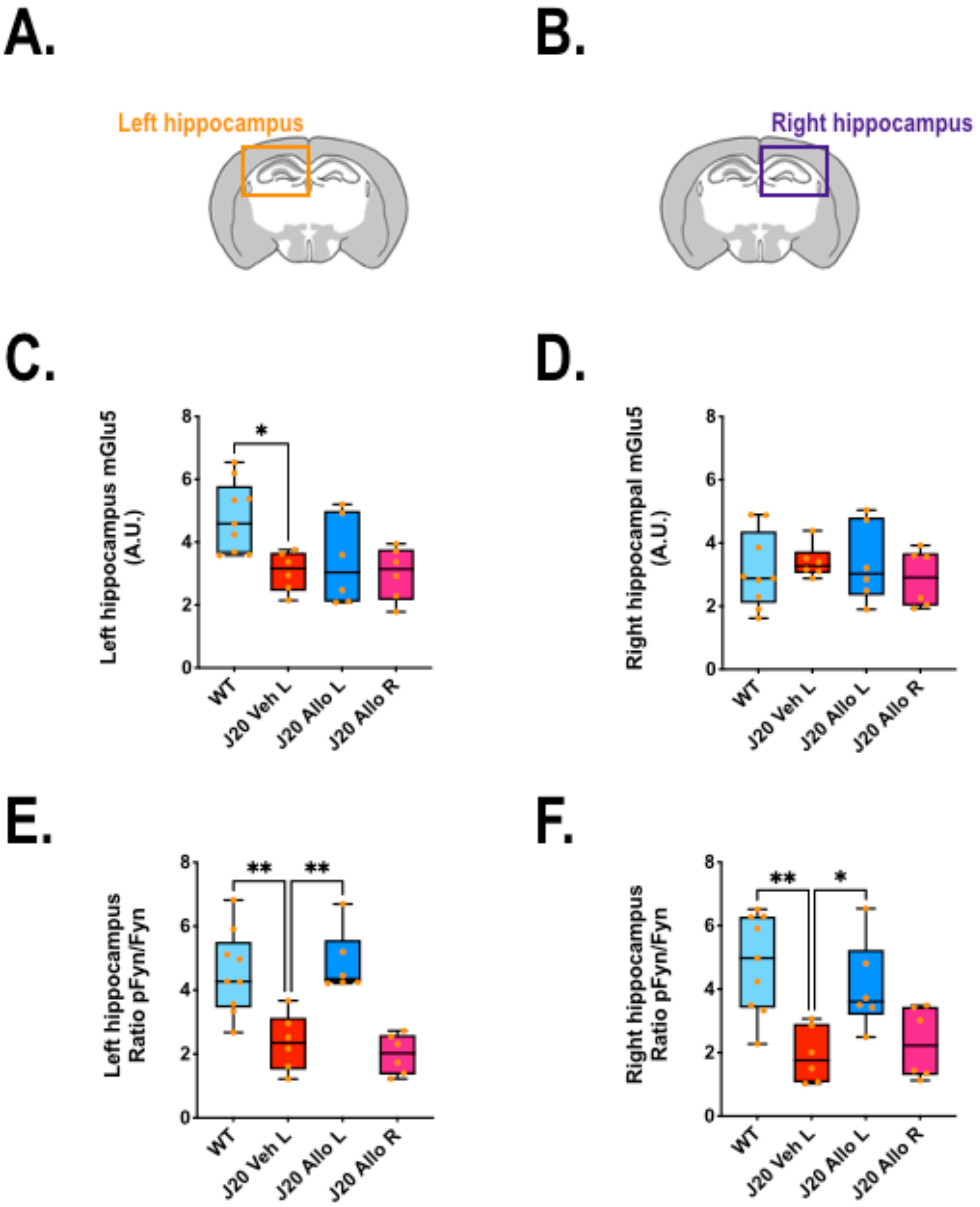

### Sup Fig.1

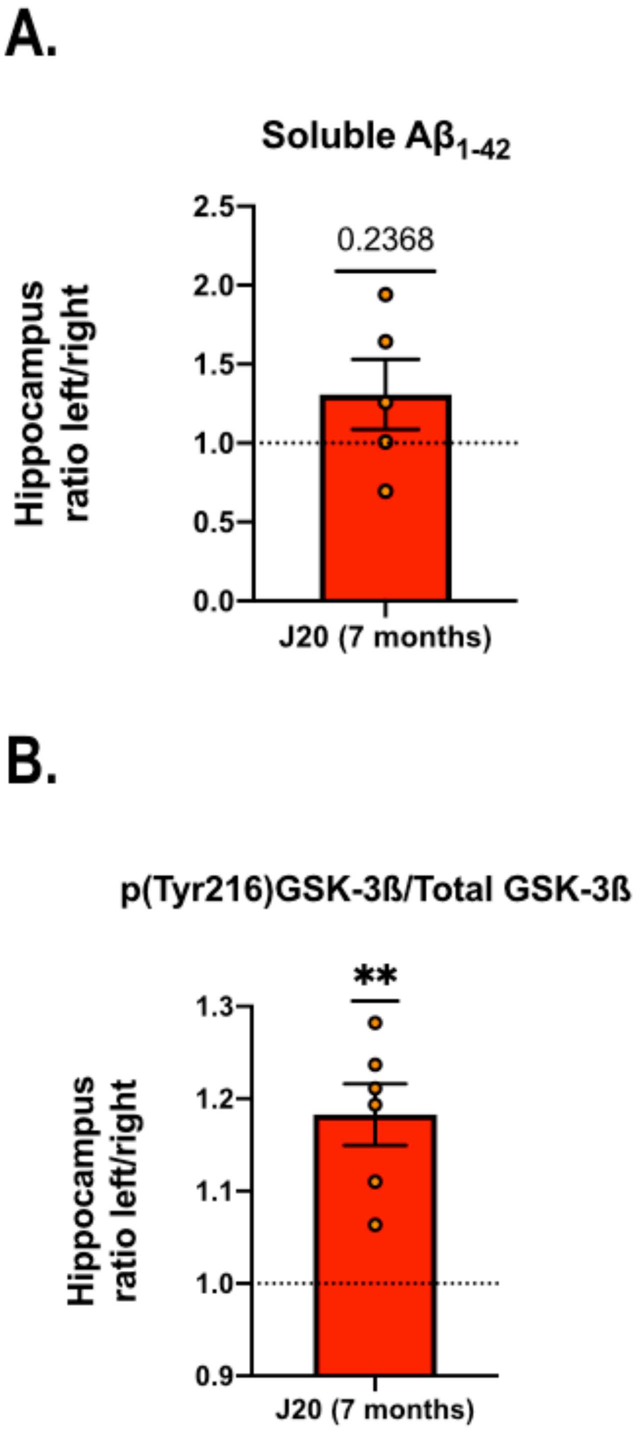
